# Functional Pairing of TCR-peptide-MHC Interactomes by Single-cell Clonal Expansion

**DOI:** 10.64898/2026.08.18.745550

**Authors:** Liu Daisy Liu, Seung Won Shin, Kevin Joslin, Chunyu Wang, James Pai, Rong Ma, Xinyu Xiang, Iain C. Clark, K. Christopher Garcia

## Abstract

T cell antigen-specific immunity depends on pairwise interactions between T cell receptors and peptide-MHC, yet isolating the TCR-pMHC pairs that drive productive engagement remains a major obstacle for antigen-specific therapeutics and for decoding TCR specificity. We overcome this by co-encoding TCR and pMHC in a single founder cell, then clonally expanding it inside a semi-permeable capsule so that genetically identical daughter cells engage in trans. T cell activation, rather than binding affinity, is used to sort cells with functional pairs, and a single PCR on the clone’s linked genomic library captures both partners. This platform, LINC-seq, recovered known cognate pairs from pooled libraries at up to 95% accuracy and performed simultaneous, library-on-library deep mutational scanning of both partners. Wild-type clonotypes ranked among the top-enriched sequences in complex mixtures, and the screens resolved co-evolutionary epistasis and cross-reactivity rules inaccessible to one-sided mutagenesis. The approach generalizes to any receptor-ligand pair whose trans-engagement drives a reporter.

## Introduction

Much of the information that organizes a multicellular organism is mediated by the pairwise interaction between cell-surface receptors and the cognate ligands they encounter on neighboring cells. These interactions define immune and neural synapses, developmental fate, and inter-organ cues, yet the functional cell-surface interactome remains incomplete: thousands of receptors and ligands lack known partners^1,2^. Even when partners are known, the rules by which sequence variation tunes productive engagement, i.e. how extracellular interactions shape intracellular signaling, remain largely speculative^1–4^.

The T cell receptor (TCR)–peptide-MHC (pMHC) system is a prime example: productive receptor-ligand engagement underlies adaptive immunity and drives pathogen defense, autoimmune disease, and anti-tumor immunity^5–7^. Yet for most TCRs, the pMHC ligands they recognize are not known. For example, the antigens recognized by the vast majority of tumor-infiltrating TCRs remain orphaned, even though this interaction lies at the center of both natural and engineered T cell responses to cancer^8,9^. What makes this particularly challenging is the immense sequence diversity present in both partners. In addition, like many protein interactions, TCR-pMHC affinity alone does not predict function. TCR–pMHC interactions are transient and low-affinity (K_D_ ∼ 1–100 μM), and many pMHC molecules that bind a given TCR with measurable affinity fail to trigger signaling, while weaker binders can be potent agonists^10–12^. Binding-based screening methods are therefore unreliable predictors of productive receptor-ligand engagement. Decoding TCR specificity requires library-on-library functional mapping in which both the receptor and the ligand are varied simultaneously^13^.

Solving this problem requires a functional assay of cell–cell interactions that meets four requirements in sequence: 1- cells must be paired at high-throughput, 2- they must remain engaged long enough for downstream signaling to occur, 3- the signal must be converted into a stable record, and 4- the molecular identities of both partners must be co-recovered. No existing method satisfies all four of these constraints at once. Soluble protein–protein binding assays, such as pMHC tetramer staining and yeast surface display, scale to library size but report only binding, missing low-affinity interactions and yielding binding interactions that do not activate the TCR^12,14,15^. Viral entry methods^16–18^ display antigen on a virus, record recognition as a single binary entry event, and read pairing information by single-cell sequencing. These methods achieve library-on-library selections at a small scale while reducing the interaction to recognition alone, without the broader cellular context or co-receptor engagement at the immune synapse. Bulk co-culture activation assays preserve this biology but lose pairing information when the transiently bound, mechanically unstable doublet dissociates. New encapsulation technologies such as open nanovials^19^ protect doublets through sorting but remain strongly weighted toward adhesion strength, missing transient interactions that still strongly activate the TCR. Proximity labeling approaches can record such cell-cell contacts and are well suited for single-sided library screens^20^. Recent droplet methods co-encapsulate a T cell with an antigen-presenting cell to preserve both the biology and pairing record but rely on single-cell sequencing to recover pairing, limiting scale^21,22^. These are all valuable approaches, yet none is capable of fully capturing the receptor-ligand epistasis that shapes functional recognition.

We developed a high-throughput cell-based functional screening platform, named LINC-seq (**L**inking **I**nteracting **N**eighboring **C**lones by **S**equencing), that sequences the paired identities of T cell receptors and peptide-MHC ligands involved in a productive engagement. The TCR–pMHC pairing problem is a particularly stringent test of any cell–cell interaction platform because the functional output is dominated by epistasis across both partners. Single substitutions on either the receptor or the peptide can switch productive engagement to non-productive engagement, and only joint covariation across both interfaces can map the underlying rules. We show that the platform recovers known cognate TCR–pMHC pairs and reveals co-evolutionary rules invisible to one-sided mutagenesis. This work, to our knowledge, is the first deep mutational scan of a TCR–pMHC interface that selects on productive T cell signaling rather than *in vitro* binding affinity. Conceptually, the approach is general: any cell-surface receptor–ligand pair whose engagement drives a transcriptional output can be screened by the same logic.

## Results

### Capturing functional receptor–ligand engagement between cell pairs

We developed a cell-based screening platform, LINC-seq (**L**inking **I**nteracting **N**eighboring **C**lones by Sequencing), to map ligand-receptor interactions that result in functional cell-cell interactions at library scale (**Figure 1A**). The method captures TCR-pMHC interactions in their native cellular context, isolates interacting cells using a reporter, and maintains genetic linkage among library variants through sorting and sequencing. The core idea is to encode both receptor and ligand within a single cell in cis, then expand that founder cell clonally inside an isolated, semi-permeable capsule so that genetically identical daughter cells engage one another in trans (**Figure 1B**). This design has several important advantages for cell-based screens. First, because the receptor-ligand pair is encoded within the same cell, libraries can be constructed and handled as single cells in bulk, eliminating the need to maintain fragile cell–cell doublets during handling and sorting. Low-affinity, transient interactions, such as the fast-off-rate engagement between a TCR and its stimulating pMHC, are captured. Second, the physical linkage of receptor and ligand sequences within the same genomic molecule allows a single PCR to directly recover the pairing information, without the need for barcoding. Third, clonal expansion creates pairs in every cell-containing capsule, reducing throughput loss due to random co-encapsulation, which limits pairing efficiency to ∼1%. Together, these features make functional receptor–ligand screening practical at a library-on-library scale.

**Figure 1.**
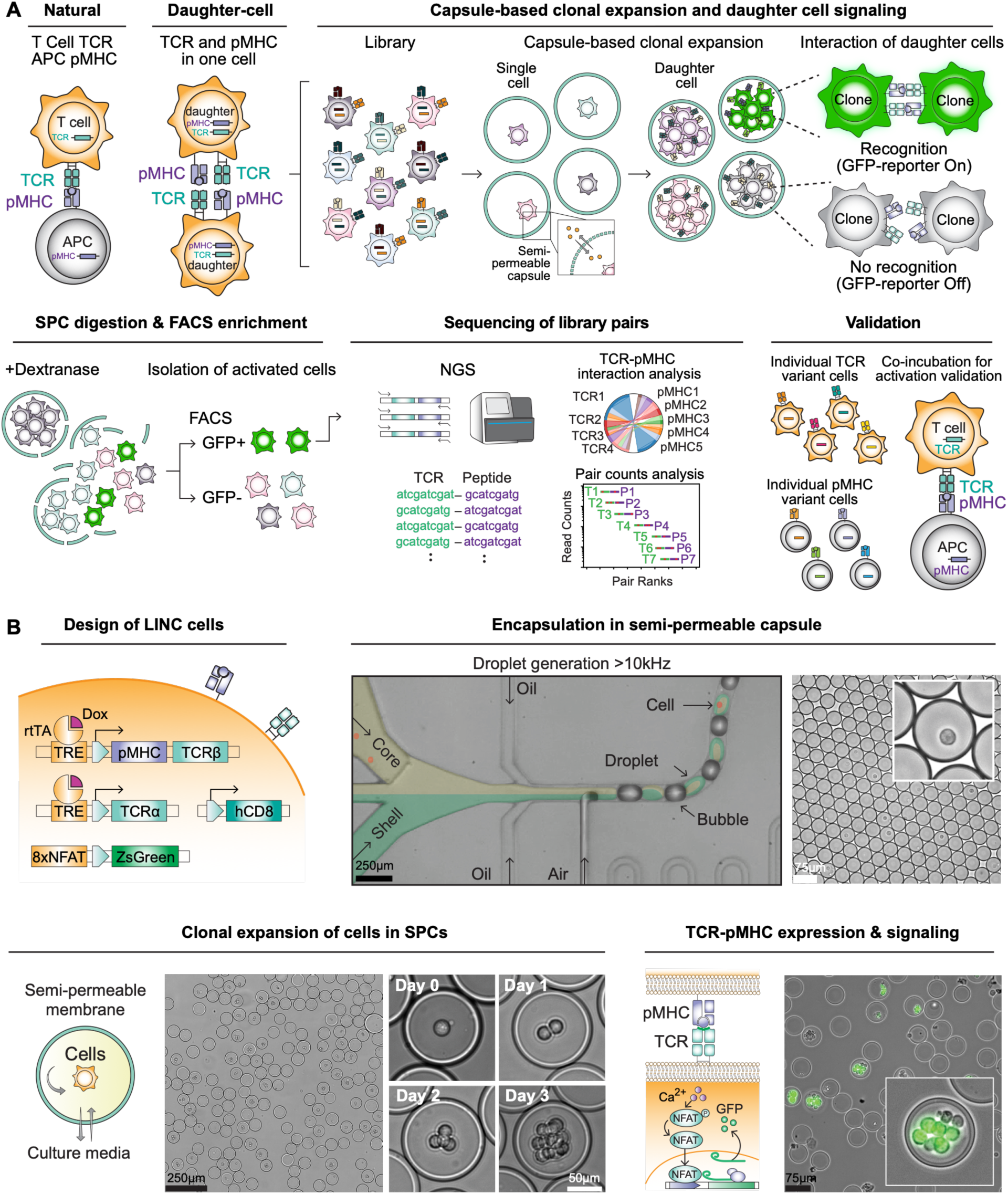
Capturing functional TCR–pMHC engagement between cell pairs. **(A)** Overview of the daughter-cell signaling strategy for detecting cell–cell receptor–ligand engagement. In conventional T cell–APC recognition, TCR and pMHC are expressed on opposing cells. In the daughter-cell format, TCR and pMHC are encoded within the same founder cell and functionally tested in trans after clonal expansion, allowing neighboring daughter cells within the same capsule to interact through matched TCR–pMHC pairs. A pooled library of TCR and pMHC variants is encapsulated into SPCs, where single founder cells clonally expand into multi-cell colonies. Cognate TCR–pMHC engagement between daughter cells activates the NFAT reporter, resulting in ZsGreen1 expression. Capsules are digested with dextranase, ZsGreen1⁺ cells are sorted by FACS, and paired TCR–pMHC sequences are recovered by next-generation sequencing. Candidate interactions are subsequently validated using individual TCR and pMHC variant cells or conventional co-incubation assays. **(B)** Design and implementation of LINC reporter cells in SPCs. LINC cells contain Dox-inducible TCR and pMHC expression modules together with an NFAT-driven ZsGreen1 reporter. Cells are encapsulated in semi-permeable capsules generated by a jet-triggered microfluidic device, followed by clonal expansion from a single founder cell over three days. The semi-permeable membrane supports media exchange while retaining cells inside each capsule. Upon doxycycline induction, TCR and pMHC are expressed on daughter cells within the same clonal colony, enabling TCR–pMHC engagement, NFAT activation, and ZsGreen1 reporter expression. Representative brightfield and fluorescence images show capsule generation, clonal colony formation, and reporter activation in SPCs. Scale bars, as indicated.

### Development of trans-signaling daughter cells

To capture productive TCR–pMHC signaling in daughter cells after clonal expansion, we engineered a single cell to encode three elements: the pMHC ligand, the TCR, and a reporter of productive interaction. TCR activation reporters typically couple a TCR-responsive regulatory element to a minimal promoter that drives a payload gene, such as a fluorescent protein (**Figure 2A**). To identify the configuration with the highest signal-to-noise, we screened a panel of TCR-responsive regulatory elements, including varying copy numbers of nuclear factor of activated T cells (NFAT) binding repeats^23^, synthetic NR4A motifs^24^, and other synthetic motifs^24^ paired with an IL-2 minimal promoter (IL2miniP), CMV minimal promoter (CMVminiP), or synthetic minimal promoter (SMP)^24^ driving mCherry expression. Reporters were individually transduced into Jurkat T cells expressing the YLQ7^25^, A6C134^26^, or DMF5 TCR^27^, and stimulated with K562 antigen-presenting cells overexpressing cognate peptides in single-chain trimer (SCT) format. Among all configurations tested, the 12×NR4A-SMP element produced a 9-fold induction in YLQ7 Jurkat cells (**Figure 2B**). Reporters built on NFAT repeats outperformed the NR4A configurations, with reporter output scaling as a function of repeat copy number. The 8×NFAT-CMVminiP configuration yielded the highest signal-to-noise ratio across all three TCRs (**Figures S1A and S1B**), reaching up to a 67-fold induction in YLQ7 Jurkat cells (**Figure 2B**). This configuration was selected for all subsequent experiments.

**Figure 2.**
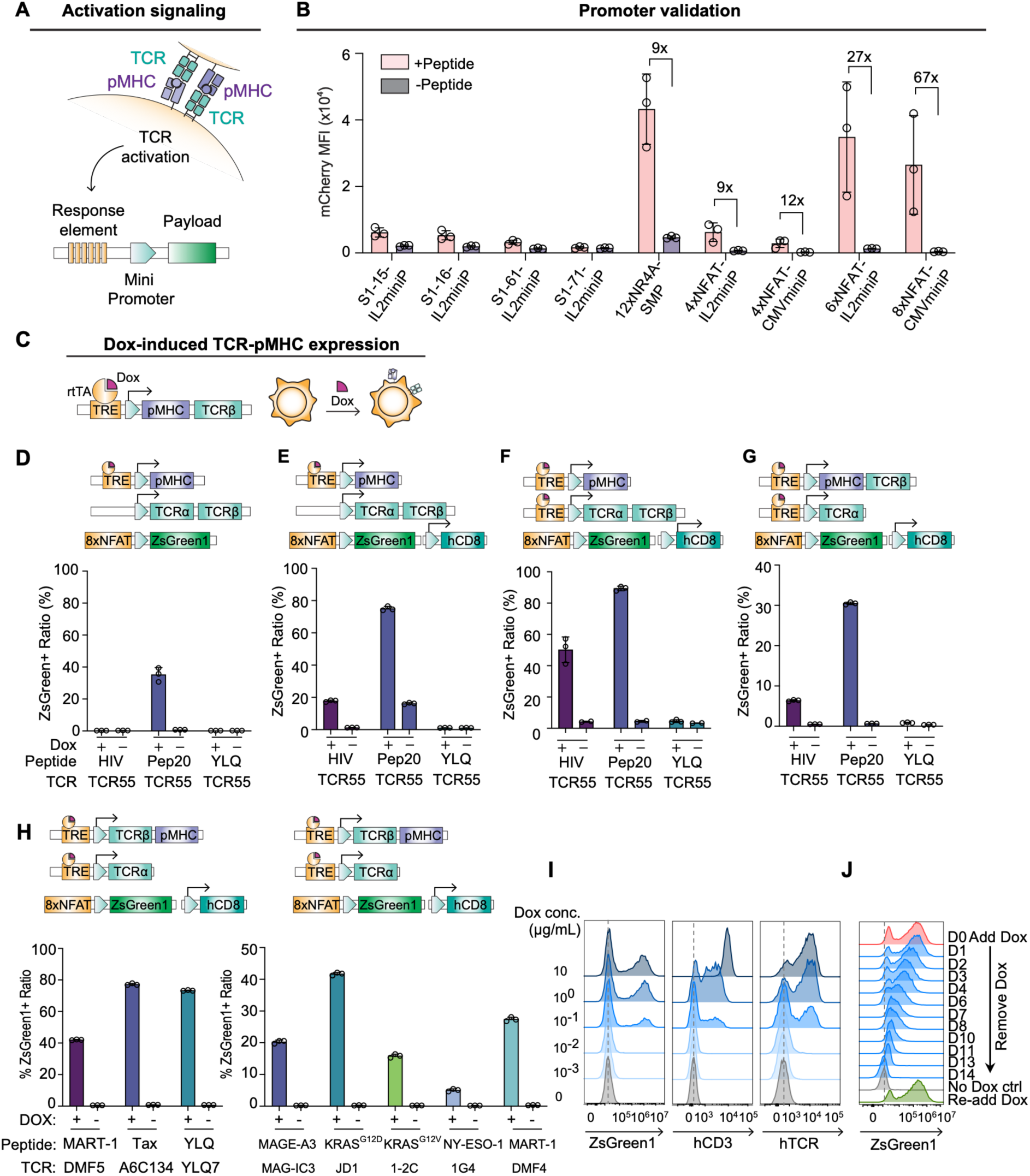
Development of the trans-signaling daughter cells. **(A)** Schematic of TCR-driven activation signaling and reporter design. Trans engagement of TCR and pMHC triggers TCR activation, which drives a response element placed upstream of a minimal promoter to express a downstream payload. **(B)** Promoter validation. Candidate response elements driving mCherry were screened in YLQ7 TCR- expressing Jurkat T cells co-cultured with antigen-presenting cells expressing the YLQ peptide as a single- chain trimer (SCT). Bars show mCherry mean fluorescence intensity (MFI) with (+Peptide) or without (– Peptide) cognate peptide; fold induction over the no peptide baseline is indicated above selected bars. **(C)** Construct schematic of Dox-inducible pMHC–TCRβ expression, in which rtTA drives transcription from a TRE promoter upon Dox addition. **(D–E)** 8*×*NFAT-ZsGreen1 reporter cells with constitutive TCR expression and Dox-inducible pMHC were tested against moderate agonist (Pep20), weak agonist (HIV), and non-cognate (YLQ) peptides, without **(D)** or with **(E)** hCD8 co-expression. Schematics indicate construct architecture for each condition. **(F-G)** 8*×*NFAT-ZsGreen1 reporter cells co-expressing hCD8 were tested against the same three peptides: Pep20, HIV, and YLQ peptides, with dox-inducible TCR and pMHC encoded on separate constructs **(F)** or with TCRβ chain and pMHC co-encoded on a single construct **(G)**. Schematics indicate construct architecture for each condition. **(H)** Reporter cells co-expressing hCD8 and a single-construct TCRβ–pMHC were validated across additional TCR–peptide pairs: DMF5/MART-1, A6C134/Tax, YLQ7/YLQ (left) and MAG-IC3/MAGE- A3, JD1/KRAS-G12D, 1-2C/KRAS-G12V, 1G4/NY-ESO-1, and DMF4/MART-1 (right). **(I)** Flow cytometry histograms of ZsGreen1, hCD3 and hTCRα/β expression in reporter cells carrying the single-construct TCRβ–pMHC across a Dox titration (0–10 µg/mL), showing dose-dependent induction that ZsGreen1 upregulation saturates at 1 µg/mL. **(J)** Flow cytometry histograms of ZsGreen1 expression monitored over 14 days following Dox removal and after Dox re-addition at day 14. A no-Dox control is shown. Data in (B), (D)-(H) are presented as mean ± SD; n = 3 independent biological replicates.

Next, we systematically evaluated multiple lentiviral TCR and pMHC co-expression architectures (**Figures 2C-H** **and Figures S1C-S1E**). In all configurations, a dedicated 8×NFAT-ZsGreen1 cassette reports TCR activation, and a doxycycline (Dox)-inducible TRE3G promoter^28^ controls pMHC SCT expression, preventing premature signaling during bulk culture and ensuring TCR–pMHC engagement occurs only upon Dox addition (**Figure 2C**). We first transduced reporter cells harboring one of three inducible pMHC SCTs, including a strong agonist (Pep20)^12^, a weak agonist (HIV, Po_l448–456_)^12^ and a non-cognate control (YLQ, SARS-CoV-2, S_269-277_)^29^, with a constitutive vector co-expressing the full-length TCR55 TCRα and TCRβ chains. This configuration yielded ∼40% ZsGreen1-positive cells with Pep20, with no activation against HIV or YLQ (**Figure 2D**). Incorporating hCD8 co-receptor overexpression doubled Pep20-driven activation to ∼80% and unmasked ∼20% activation against the weak agonist HIV, while also producing ∼20% Dox-independent background signal (**Figure 2E**). Placing TCR expression under Dox-inducible control eliminated this background and maintained ∼80% activation against Pep20 and ∼40% against HIV, with HIV-specific activation strictly dependent on hCD8 co-expression (**Figure 2F**, **Figure S1C**). Dox titration further revealed graded, dose-dependent induction of hCD3, TCRαβ surface expression, and ZsGreen1 reporter signal, saturating at 100 ng/ml (**Figure S1F, S1G**).

Library-on-library screening depends on recovering the identity of TCR–pMHC pairs from each productive engagement. Clonal expansion, trans signaling between daughter cells, and NFAT-based sorting remove the need to isolate cell pairs, but the genetic pairing itself must still be recovered at single-cell resolution from the sorted population. When library components are encoded at separate genomic loci, recovering their linkage by sequencing can be accomplished with single-cell barcoding. However, single-cell sequencing after sorting limits the scalability of screens and increases the experimental complexity. To avoid this, we co-express the pMHC SCT and TCRβ chain on a single inducible vector via a P2A self-cleaving sequence. This physical linkage on a single lentiviral construct allows a single PCR to recover both the pMHC and TCR identity, removing the need to barcode separate molecules. The linked architecture produced ∼30% ZsGreen1 positive cells against Pep20 and ∼10% against HIV in a hCD8-dependent manner, with no activation against non-cognate YLQ (**Figures 2G**, **S1D and S1E**). Inverting the construct to place TCRβ upstream of the pMHC SCT modestly improved activation in A6C134 (64% to 76%) and YLQ7 (61% to 72%) reporter cells (**Figure 2H**), and this optimized construct was validated across eight additional TCR–pMHC pairs including DMF5 TCR/MART-1/HLA-A*02^27^, A6C134 TCR/Tax/HLA- A*02^26^, YLQ7 TCR/YLQ/HLA-A*02^25^, MAG-IC3 TCR/MAGE-A3/HLA-A*01^30^, JD1 TCR/KRAS- G12D/HLA-A*11^31^, 1-2C TCR/KRAS-G12V/HLA-A*11^32^, 1G4 TCR/NY-ESO-1/HLA-A*02^33^, and DMF4 TCR/MART-1/HLA-A*02^27^, confirming broad compatibility across different HLA allotypes and antigen systems (**Figure 2H**). The linked TCRβ–pMHC construct showed dose-dependent induction of TCRαβ surface expression and ZsGreen1 reporter signal, saturating at 1 μg/ml (**Figure 2I**). ZsGreen1 expression returned to baseline within 14 days of Dox removal and was restored upon Dox reintroduction (**Figure 2J**). The resulting reporter cell co-expresses a P2A-linked TCRβ and pMHC single-chain trimer from a single Dox-inducible vector and reads out productive engagement through an 8×NFAT-ZsGreen1 cassette, resolving agonists across a range of potencies in a fully inducible, reversible manner. This cell is the unit that LINC-seq encapsulates, expands into daughter cells, and screens for productive engagement.

### Capsule-based daughter cell signaling

To enable daughter cell signaling, a single founder cell must expand clonally into a colony of genetically identical cells that remain viable, signaling-competent, and independent of other colonies. We first attempted to use water-in-oil droplets for this purpose but observed poor growth and limited reporter signaling (**Figure S2**), likely because small (50-300 pL) water-in-oil droplets become nutrient-depleted within hours. We therefore explored semi-permeable capsules formed by aqueous two-phase separation^34–37^. Capsules are generated by co-flowing immiscible aqueous polymers in droplets using microfluidics (**Figure 1B**). The aqueous phases separate into a core and a shell, and the shell is polymerized to form semi-permeable capsules (SPCs) that maintain cellular compartmentalization while allowing ongoing media exchange during multi-day culture. For cell culture applications, capsule formation requires compatibility of aqueous phase separation and shell crosslinking with cell partitioning, viability, and recovery. We evaluated several Polyethylene glycol (PEG)-based ATPS-based capsule chemistries, including Dextran- PEG-Maleimide^36^, Dextran-PEG-Ortho-Pyridyldisulfide (OPSS)^36^, and Dextran-Polyethylene Glycol Diacrylate (PEGDA)^37^, all of which had limitations related to cell compatibility or recovery (**Figure S3**).

Dextran offered an attractive basis for capsule optimization because it is a biocompatible, chemically versatile polysaccharide whose hydroxyl-rich backbone can be modified to tune solubility, hydrophobicity, and crosslinking behavior^38^. We evaluated a dextran-based capsule chemistry in which a methacrylate- and butyrate-functionalized dextran phase forms a photocurable shell around a dextran-rich, cell-containing core. We synthesized the shell polymer, confirmed methacrylate and butyrate resonances by ¹H NMR (**Figure S4A**), and generated stable capsules using droplet microfluidics, followed by oil removal (**Figure S4B, S4C**). The dextran-based SPCs supported robust clonal expansion (**Figure S4D**), facile handling, and rapid shell dissolution for cell recovery (**Figure S4E**). Jurkat T cells encapsulated in capsules grew for over seven days, maintained high viability, and reached 10-15 cells per capsule (**Figure 3A**).

**Figure 3.**
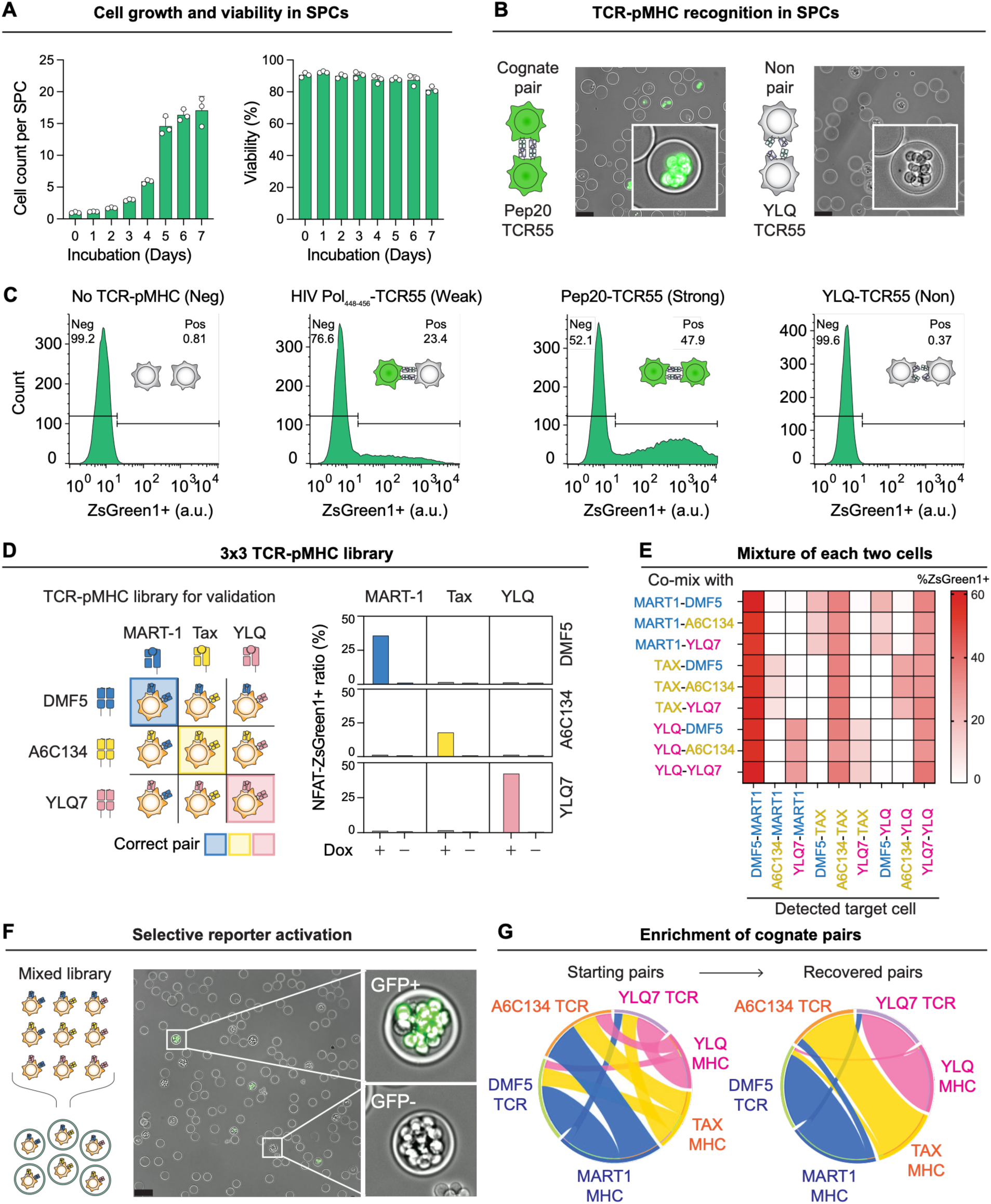
Proof-of-concept validation of the LINC-seq screening platform using a 3×3 TCR–pMHC library. **(A)** Cell growth and viability in SPCs. Cells encapsulated in SPCs expanded over seven days, while maintaining high viability throughout culture. Following encapsulation, SPCs were distributed into separate wells of a 24-well plate, and cell growth and viability were monitored independently. Bar plots show the number of cells per SPC and cell viability as a function of incubation time. Data are presented as mean ± SD; n = 3. **(B)** Representative microscopy images showing TCR–pMHC recognition in SPCs after Dox induction. Capsules containing the cognate Pep20–TCR55 pair showed NFAT-driven ZsGreen1 reporter activation, whereas capsules containing the non-cognate YLQ–TCR55 pair remained reporter-negative. Insets show representative single capsules. Scale bars, 75 μm. **(C)** Flow cytometric quantification of reporter activation across four conditions: negative control with no TCR–pMHC pair, weak cognate Pol_448-456_–TCR55, moderate cognate Pep20–TCR55, and non-cognate YLQ–TCR55. The percentage of ZsGreen1-positive cells increased according to cognate interaction strength, with the strongest activation observed for Pep20–TCR55 and minimal activation in the negative and non-cognate controls. **(D)** Validation of a 3×3 TCR–pMHC library composed of three pMHCs, MART-1, Tax, and YLQ, and three TCRs, DMF5, A6C134, and YLQ7. Only the matched cognate pairs, DMF5–MART-1, A6C134–Tax, and YLQ7–YLQ, induced Dox-dependent NFAT–ZsGreen1 reporter activation, whereas non-cognate combinations remained near background. **(E)** Pairwise co-mixing analysis of the nine TCR–pMHC cell types. The heatmap shows the percentage of reporter-positive cells detected in each target population after co-culture with each partner population, confirming selective activation by cognate intercellular TCR–pMHC engagement. **(F)** Selective reporter activation in a mixed-library SPC experiment. Following encapsulation and clonal expansion, only capsules containing productive TCR–pMHC interactions showed ZsGreen1 fluorescence, enabling activated capsules or cells to be distinguished from non-activated populations. Scale bars, 100 μm. **(G)** Enrichment of cognate TCR–pMHC pairs after reporter-based screening and sequencing. Chord diagrams compare TCR–pMHC pair distributions in the starting library and the recovered ZsGreen1 positive population, showing enrichment of the expected cognate interactions. Pairs were identified by bulk gDNA sequencing.

Next, we confirmed that cis-encoded receptors and ligands could induce TCR-dependent activation of the NFAT reporter in capsules. We built weak, strong, and non-cognate TCR-pMHC pairs and tested signaling. Inside each capsule, a single founder cell was clonally expanded into a multi-cell colony over 72 hours, so that every cell descended from the same founder and shared the same TCR–pMHC pair (**Figure 3B**). We validated specificity and dynamic range using four reporter conditions: a negative control (no TCR or pMHC), a non-cognate pair (TCR55–YLQ), a weak cognate pair (TCR55–HIV Pol_448-456_), and a moderate cognate pair (TCR55–Pep20). After Dox induction, fluorescence microscopy of SPCs showed a bright ZsGreen1 signal exclusively in capsules containing the correct cognate pairs, and an undetectable signal in capsules containing non-cognate pairs (**Figure 3B**). The proportion of ZsGreen1 positive cells as a function of Dox-treatment duration was tested to assess the temporal dynamics of reporter activation (**Figure S4F**). Flow cytometry recapitulated this hierarchy quantitatively: 47.9% ZsGreen1⁺ cells for the strong cognate pair, 23.4% for the weak cognate pair, and 0.81% and 0.37% for the negative control and non-cognate pair, respectively (**Figure 3C**). The gradient of response across cognate pairs with differing intrinsic potency establishes that the platform captures the full dynamic range of functional TCR signaling, rather than binary events.

### Proof-of-concept validation of the LINC-seq platform using a 3×3 TCR–pMHC library

To stress-test cognate-pair recovery, we manually constructed a 3×3 matrix of three TCRs and three pMHC complexes spanning well-characterized cognate pairs: DMF5–MART-1 (Melanoma Antigen Recognized by T-cells 1, HLA-A*02:01), A6C134–Tax (HTLV-1, HLA-A*02:01), and YLQ7–YLQ (SARS-CoV-2 spike, HLA-A*02:01) (**Figure 3D**, **left**). Each TCR was cloned in combination with each pMHC, generating nine distinct reporter cell lines. Bulk culture of individual cell lines following Dox induction produced a strong reporter signal exclusively on the diagonal of the matrix, with negligible off-diagonal signal (**Figure 3D**, **right**), confirming that the all-in-one reporter chassis recapitulates known TCR–pMHC specificity. Pairwise co-culture of all nine cell lines also produced the expected trans-complementation: two lines that are each non-cognate on their own reconstituted a productive interaction when mixed, because one line’s TCR recognized the other line’s pMHC (**Figure 3E**). For example, a MART-1–A6C134 line co-cultured with a Tax–YLQ7 line generated a signal (via Tax–A6C134), even though neither line could signal in isolation. This result establishes the necessity of single-cell encapsulation to prevent cross-activation and preserve pairing fidelity.

We next performed a pooled mini-screen to test whether cognate pairs could be recovered from the 9- member library. The small library was encapsulated as single cells in SPCs (∼10^5^ capsules per replicate), expanded for three days, and induced with Dox for two days to drive library expression. Cells activating the reporter were sorted for ZsGreen1⁺ signal (**Figure 3F**), and paired peptide–TCR CDR3β sequences were recovered by next-generation sequencing. Pre-sort, the nine TCR–pMHC combinations were distributed approximately uniformly, consistent with random mixing (**Figure 3G**, **left**). Post-sort, the correct pairs were sharply enriched, with >95% of reads mapping to one of the three cognate combinations (**Figure 3G**, **right**). We confirmed this recovery with two orthogonal readouts: Sanger sequencing of TCR– pMHC pairs amplified from bulk genomic DNA matched a cognate combination in 75 of 83 pairs (90% accuracy), and independent single-cell RT-PCR of sorted ZsGreen1⁺ cells reached 95% accuracy (129 of 135 pairs) (**Figure 3G**, **Figure S5**). Together, these readouts show that trans engagement within clonal SPCs reliably discriminates cognate from non-cognate pairs in a pooled, single-pot format.

### LINC-seq reveals convergence onto the wild-type sequence at the low-affinity DMF4-MART-1 interface

As our next test of the LINC-seq workflow, we attempted a large-scale library-on-library mutagenesis screen of TCR CDR3 loops, which dominate peptide recognition specificity, against a peptide library. We selected the low-affinity DMF4 TCR/MART-1/HLA-A*02 pair, in which MART-1 peptide serves as a weak agonist for DMF4 TCR^39^, and focused library design on the CDR3β loop and its cognate peptide contact residues, as revealed from a crystal structure of the complex^40^. Reporter cells stably expressing the 8*×*NFAT-ZsGreen1 reporter and hCD8 were sequentially transduced with the Dox-inducible DMF4-TCRα construct and the linked CDR3β–peptide library at low MOI, ensuring each cell harbored a single unique TCR–pMHC combination (**Figure 4A**). Crystal structure analysis of the DMF4–MART-1 complex (PDB 3QDM)^40^ identified CDR3β residues V96, G97, V98, G99, and Q100 as the primary contacts with peptide residues L8 and T9 (**Figure 4B**). A combinatorial library was designed by simultaneously randomizing these five CDR3β positions and two peptide contact positions using degenerate codons, generating an estimated diversity of approximately 1.8×10^7^ unique TCR–pMHC sequence combinations spanning the joint contact interface (**Figure 4C**). To reach the throughput required for library-scale screening, we employed a jet-triggered droplet generator in which periodic air bubbles segment the aqueous stream, producing monodisperse SPCs at ∼10 kHz, an order of magnitude faster than conventional flow-focusing systems^41,42^.

**Figure 4.**
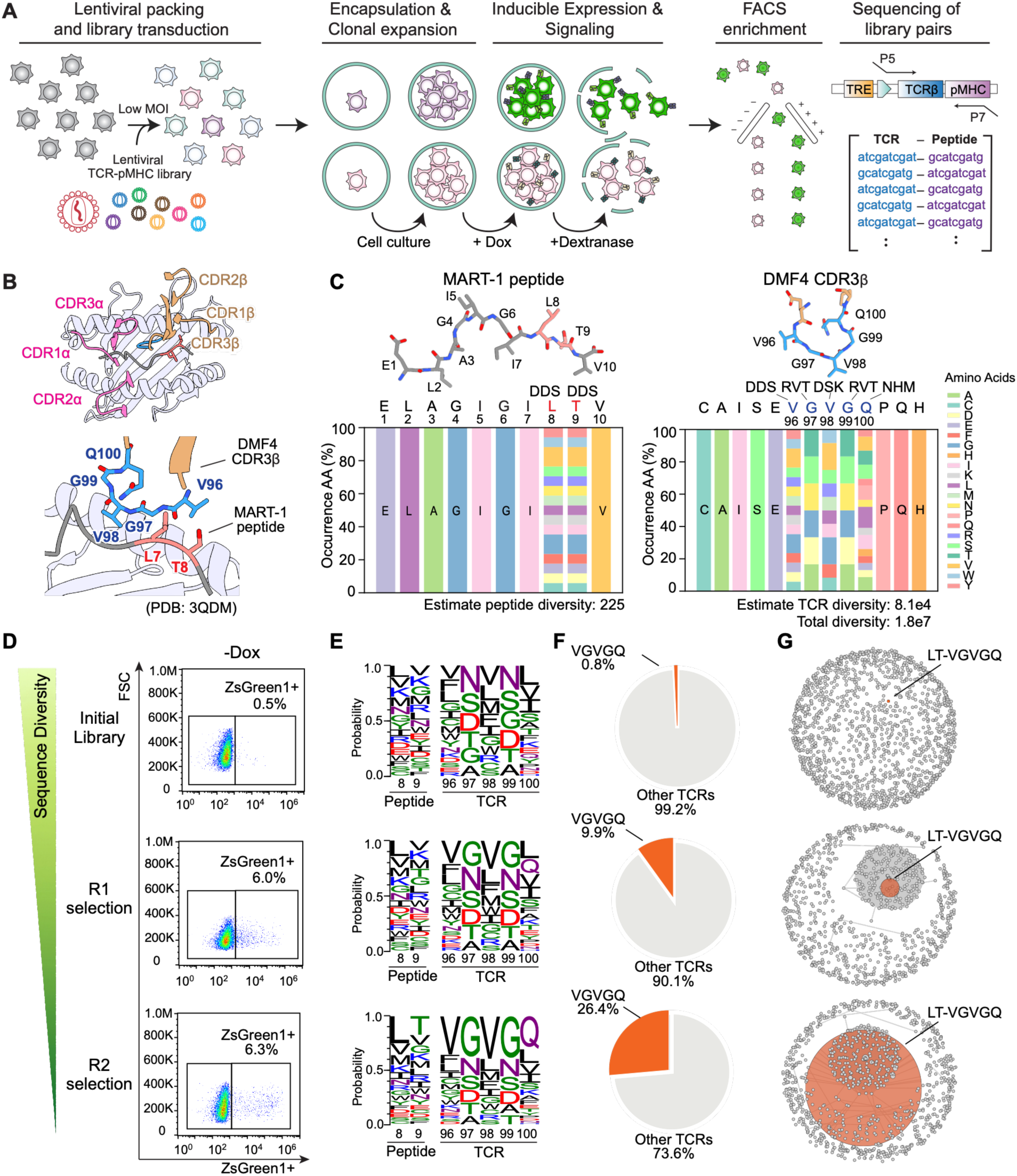
LINC-seq reveals convergence to the wild-type sequence at the low-affinity DMF4–MART- 1 interface. **(A)** Overview of the library-on-library TCR-pMHC screening with LINC-seq. Cells are transduced with a TCR–pMHC library at low MOI, encapsulated in SPCs for clonal expansion, induced with doxycycline to trigger reporter signaling, and released by dextranase treatment. Reporter-positive cells are enriched by FACS, and the linked TCR–pMHC pairs are recovered by sequencing. **(B)** Crystal structure of the DMF4–MART-1 TCR–pMHC complex (PDB: 3QDM). Top: overview with TCRα CDR loops (pink) and TCRβ CDR loops (orange) labeled. Bottom: close-up of the interface between CDR3β residues (blue) and peptide (red). **(C)** Position-frequency matrices showing the amino acid diversity introduced at each mutagenized position in the MART-1 peptide (left) and DMF4 CDR3β (right) libraries. The peptide and CDR3β contact residues are shown as sticks above each matrix, with mutagenized sites labeled in red (peptide) and blue (TCR) and the degenerate codons indicated above each position. Estimated library diversities are shown below. **(D)** Flow cytometry dot plots showing ZsGreen1-positive frequencies in library cells without DOX (baseline), after the first round of bulk activation and sorting (R1), and after a second round of SPC encapsulation, activation, and sorting (R2). The wedge indicates the progressive narrowing of sequence diversity across rounds. **(E)** Sequence logos of the mutagenized peptide (positions 8 and 9) and TCR CDR3β (positions 96–100) in the initial library and after ZsGreen1 positive enrichment at R1 and R2, showing convergence toward the wild-type sequence (LT and VGVGQ). **(F)** Pie charts showing the fraction of VGVGQ-containing TCRs in the initial library and after ZsGreen1 positive enrichment at R1 and R2. **(G)** Sequence similarity network (SSN) of the top 1000 TCR–pMHC pairs across screening rounds. Each node is a unique sequence, with node size scaled to read count, and edges drawn at an edit-distance threshold of 2. The wild-type pair (LT-VGVGQ, orange) is an isolated node in the starting library and expands into a dominant clonotype by R2.

Library cells were then processed through two sequential rounds of selection to increase screening throughput. In the first round (R1), 200 million bulk-cultured library cells were induced with Dox and MACS-enriched for T cell activation marker CD69 prior to ZsGreen1 sorting, yielding 6.0% ZsGreen1 positive cells against a 0.5% uninduced background. After a 14-day resting period to allow reporter signal to return to baseline, 10 million cells were encapsulated in SPCs and re-induced in the second round (R2), yielding 6.3% ZsGreen1-positive cells (**Figure 4D**). Paired TCR–pMHC sequences were amplified from the genomic DNA of the starting library and both (R1, R2) ZsGreen1+ populations. Sequence logo analysis of the mutagenized positions revealed progressive convergence toward wild-type amino acids at both CDR3β positions (VGVGQ) and peptide contact positions (LT) across the two selection rounds, while all randomized positions showed nearly uniform amino acid distributions in the starting library (**Figure 4E**). This convergence was confirmed by direct quantification, with VGVGQ frequency increasing from 0.8% in the starting library to 9.9% after R1 and 26.4% after R2 (**Figure 4F**). Sequence similarity network analysis of the top 1000 pairs from each sample independently revealed that the wild-type DMF4–MART- 1 pair was a rare, isolated node in the starting library but progressively expanded into a dominant clone by the second round, and wild-type DMF4 TCRs formed a dense cluster across the selection (**Figure 4G**). Together, these results demonstrate that SPC-based selection accurately recovers functionally active cognate pairs from a large combinatorial library and progressively enriches dominant sequences across successive rounds. The pronounced and selective enrichment of the wild-type pair reflects its strong competitive advantage within the mutagenized sequence space and suggests that further optimization of this weak agonist interaction may require engineering CDR regions beyond CDR3β.

### A broadened landscape of functional variants at the moderate-affinity TCR55-Pep20 interface

We next applied the platform to a moderate-affinity TCR-pMHC pair presented by a different HLA allotype, mapping its full coevolutionary sequence landscape. The Pep20 peptide presented on HLA-B*35 serves as a high-potency agonist ligand for the TCR55 TCR^12^. The crystal structure of TCR55–Pep20 complex (PDB: 6BJ8) revealed the CDR3β residues R96, G97, G98, and T99 as the primary contacts with Pep20 peptide residues E5 and A7 (**Figure 5A**). A combinatorial library carrying degenerate codons at four CDR3β positions and two peptide contact positions was generated, with an estimated diversity of ∼6.8×10^6^ unique TCR–pMHC pairs (**Figure 5B**).

**Figure 5.**
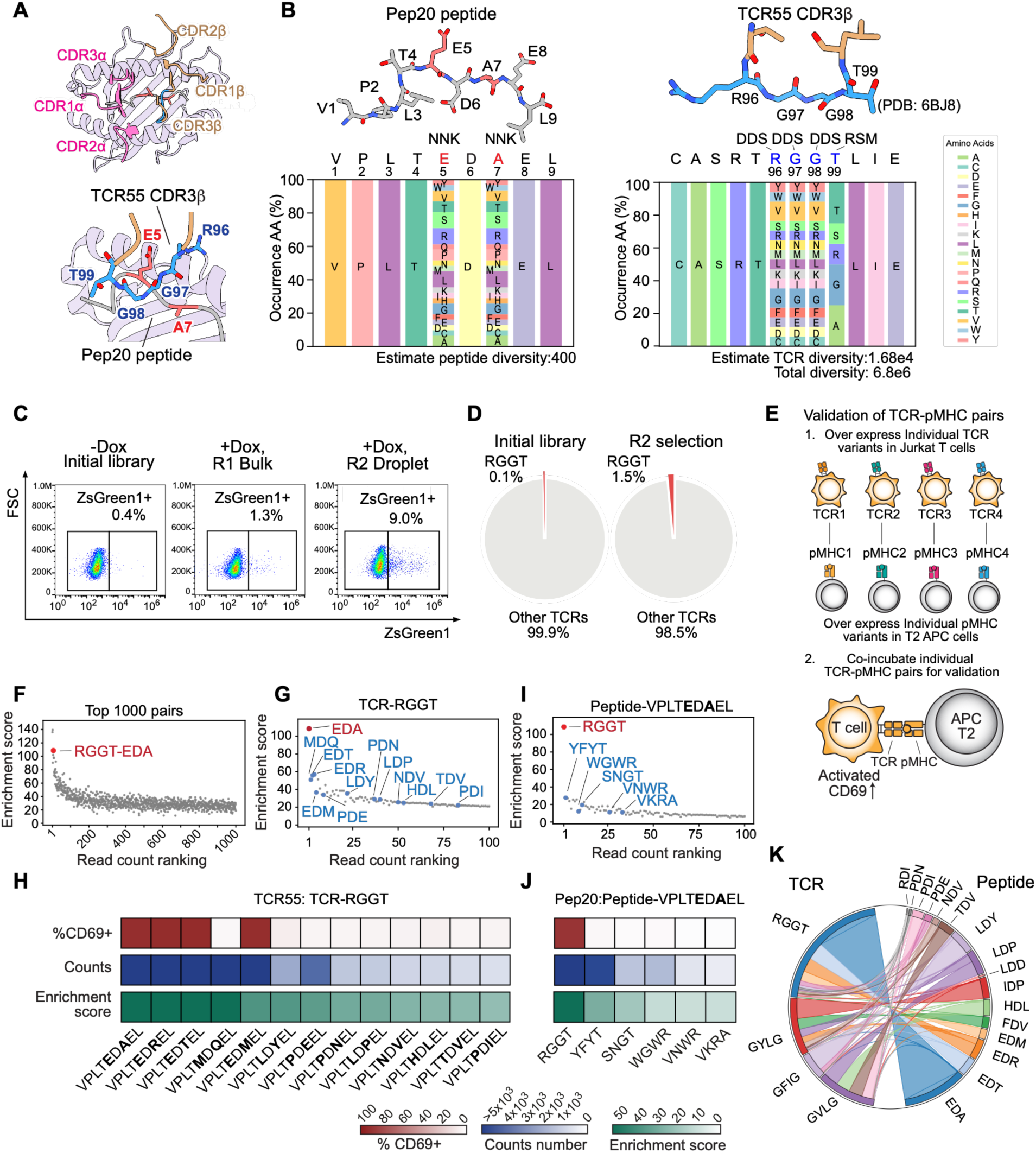
LINC-seq reveals a broadened functional landscape at the moderate-affinity TCR55–Pep20 interface. **(A)** Crystal structure of the TCR55–Pep20 TCR–pMHC complex (PDB: 6BJ8). Top view: TCRα and TCRβ CDR loops are shown in pink and orange, respectively. Bottom view: the CDR3β–peptide interface, with CDR3β in blue and pep20 peptide in red. **(B)** Position-frequency matrices showing amino acid diversity introduced at each mutagenized position in the peptide (left) and CDR3β (right) libraries. Degenerate codons are indicated above each position, and peptide and TCR CDR3β contact residues are shown as sticks above each matrix. Estimated library diversities are shown below. **(C)** Flow cytometry dot plots showing ZsGreen1-positive frequencies in library cells without Dox (baseline), after first round of bulk activation and sorting (R1), and after a second round of droplet encapsulation, activation, and sorting (R2). **(D)** Pie charts showing the fraction of RGGT CDR3β clonotypes in the initial library, and after droplet-based sorting (R2). **(E)** Schematic of TCR–pMHC pair validation. Individual TCR variants are overexpressed in Jurkat T cells and cognate pMHC variants are overexpressed in T2 APC cells. TCR-expressing T cells and pMHC- expressing APCs are co-incubated as matched pairs, and T cell activation is assessed by upregulation of surface CD69. **(F)** Scatter plot showing enrichment score (log ratio of normalized read counts between the final and initial libraries divided by its standard error, y-axis, see Methods) for each of the top 1000 enriched TCR-peptide pairs, ranked by read count (x-axis). The wildtype TCR55–Pep20 pair is highlighted in red. **(G)** Scatter plot showing peptide variants paired with TCR-RGGT, with enrichment score (y-axis) for each peptide ranked by read count (x-axis). Wildtype peptide EDA is highlighted in red, other experimentally tested pairs shown in blue. **(H)** Heatmaps showing %CD69+ activation (top row), R2 sequencing read counts (middle row), and enrichment scores (bottom row) for experimental validated peptide variants paired with TCR-RGGT. Each column represents a distinct peptide as labeled below. **(I)** Scatter plot showing TCR partners of wildtype Pep20 peptide (VPLTEDAEL). Wildtype pairs are in red; other experimentally tested pairs in blue. **(J)** Heatmaps showing %CD69+ activation (top row), R2 sequencing read counts (middle row), and enrichment scores (bottom row) for TCR partners of the wildtype Pep20 peptide. **(K)** Circos plot of experimentally validated TCR-peptides pairs. Ribbons connect each TCR clonotype (left) to its paired peptide partners (right), with ribbon width proportional to read count, illustrating the breadth and relative abundance of peptide partners recovered for each TCR clonotype.

Screening was carried out using the two-round sequential procedure established for the DMF4–MART-1 screen, yielding 1.3% ZsGreen1 positive cells after bulk induction (R1) and 9.0% after SPC-based re-induction (R2), against a 0.4% uninduced background (**Figure 5C**). Paired sequences from the starting library and both sorted populations were sequenced to reveal positional enrichment at each selected CDR3β and peptide position. Among the top-ranked clonotypes recovered from the SPCs enriched ZsGreen1 positive cells, the wild-type TCR55 CDR3β (RGGT) ranked first among TCR clonotypes (**Figure S6A**). Unlike the dominant enrichment of VGVGQ observed in the DMF4–MART-1 screening, the RGGT motif showed more modest but progressive enrichment, rising from 0.1% in the starting library to 1.5% after the second round, consistent with a broader functional sequence space at this moderate-affinity interface (**Figure 5D**). Sequence logo analysis of the mutagenesis positions across samples revealed relatively uniform amino acids across the positions (**Figure S6B**).

To evaluate the activity of recovered pairs, we individually expressed a series of TCRs in Jurkat cells and the corresponding target peptides as single-chain pMHC trimers on T2 cells, then co-cultured each pair to assess T cell activation by CD69 upregulation (**Figure 5E**). The screen identified a broad set of functional TCR–pMHC pairs within the TCR55-Pep20 library. Pair enrichment analysis, using each pair’s normalized enrichment score, which weights its log fold change by the confidence of that estimate, showed that the wild-type TCR55–Pep20 pair (RGGT-EDA) ranked third among all recovered pairs, with pair frequencies declining gradually by rank (**Figure 5F**). Functional validation of the top-ranked peptide binders against TCR-RGGT showed that 4 of 5 tested peptides induced over 90% CD69 upregulation, while pairs with low read counts and low enrichment scores showed no detectable activation, confirming that a combined measure of read count and enrichment score is broadly predictive of functional activity (**Figure 5G, 5H**). When TCR sequences were instead ranked against the wild-type Pep20 peptide, RGGT appeared as the dominant clonotype while all other tested TCR binders showed no activation against wild-type Pep20, indicating an asymmetry between peptide and TCR sequence tolerance at this interface (**Figure 5I, 5J**).

To probe the breadth of functional variants further, three additional top-ranked TCR clonotypes, GYLG, GVLG, and GFIG were validated against a common panel of peptide partners spanning a range of enrichment ranks (**Figure 5K**, **S6A, S6C-E**). GVLG showed the broadest reactivity, with its four top-ranked peptides all inducing CD69 upregulation (**Figure S6C**). GYLG activated against 2 of 5 tested top-ranked peptides, and GFIG showed strong activation against its top-ranked peptides LDY (**Figure S6D, S6E**). Validation against the novel peptide VPLTLDPEL identified GVLG as a top-ranked binder, which showed significant activation, whereas the SSIT variant and lower-ranked binders showed no detectable activation (**Figure S6F**). Overall, a combined measure of read count and enrichment score was predictive of functional activation across the majority of tested pairs, with occasional exceptions in both directions. Some high-count pairs showed weak activation, such as RGGT–MQ and GFIG–PDE, whereas some low-count pairs retained strong activation, such as GVLG–IDP and GFIG–IDP, reflecting the imperfect correspondence between sequencing enrichment and functional potency. Collectively, these results demonstrate that the LINC-seq platform can discover genuinely novel high-activity TCR variants beyond the wild-type sequence, including variants that would not have been identified by rational design.

### An expansive coevolutionary landscape at the high-affinity A6C134-Tax interface

We next screened a high-affinity TCR–pMHC pair to map the coevolutionary sequence landscape at a well-characterized structural interface. The A6C134 TCR recognizes the HTLV-1-derived Tax peptide (LLFGYPVYV) presented on HLA-A*02 with a binding affinity of 4 nM^26^. Crystal structure analysis of the A6C134–Tax complex (PDB: 4FTV) identified CDR3β residues L98, M99, S100, and A101 as the primary contacts with Tax peptide residues V7 and Y8 (**Figure 6A**). Randomizing these four CDR3β positions together with the two peptide contact positions generated a paired library of ∼7.3×10^6^ unique TCR–pMHC combinations (**Figure S7A**). In the first round of screening (R1), 300 million library cells were encapsulated at three cells per capsule and induced with Dox, yielding 1.62% ZsGreen1-positive cells compared to a 0.1% uninduced background (**Figure 6B**). Enriched cells were re-encapsulated as single cells per capsule (R2), further increasing the ZsGreen1-positive cells to 6.32% upon Dox induction (**Figure 6B**). Sequence logo analysis showed modest enrichment of specific residues at both peptide and CDR3β positions across selection rounds, starting from the broad diversity of the initial library (**Figure S7B**).

**Figure 6.**
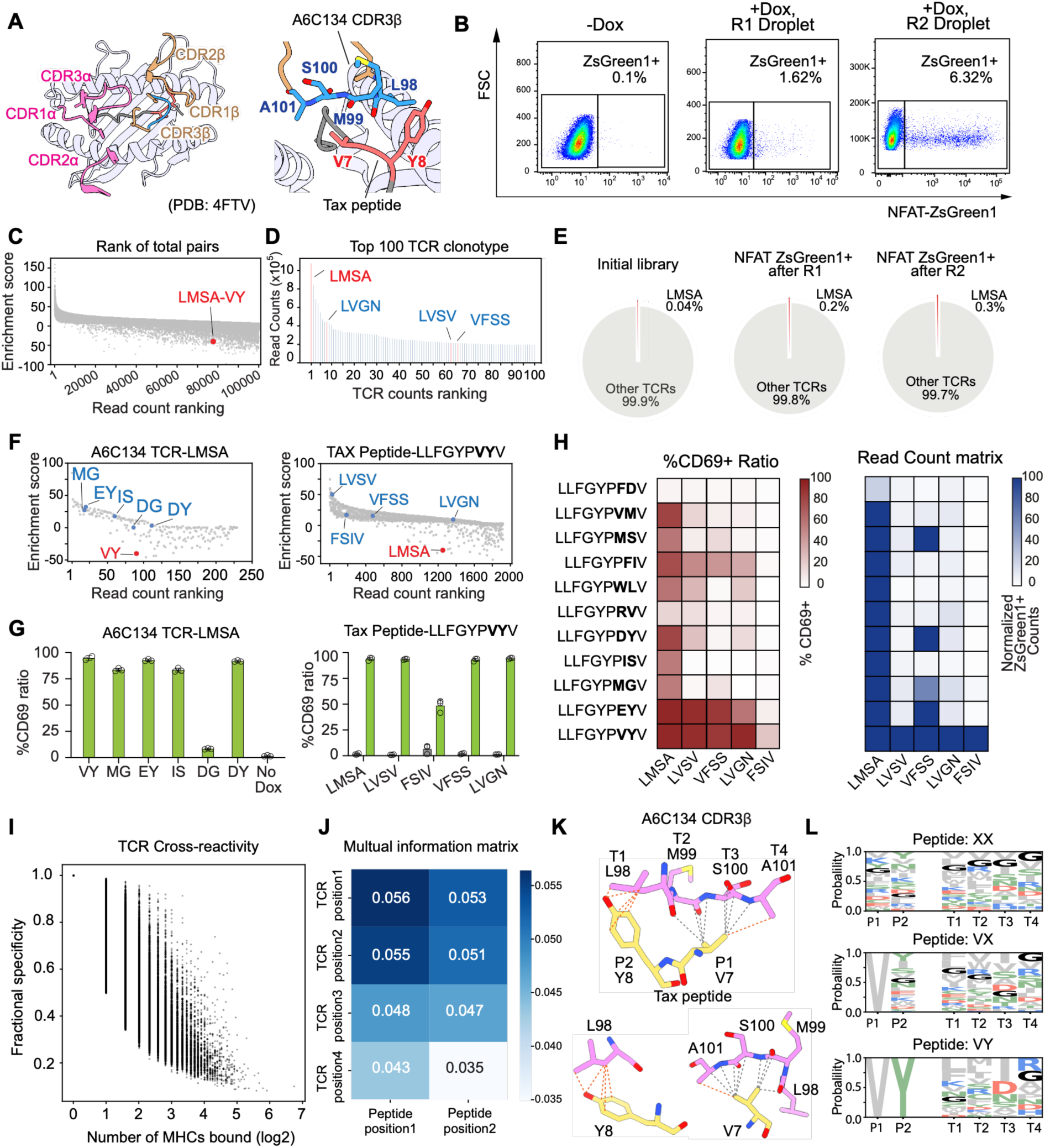
LINC-seq reveals an expansive coevolutionary landscape at the high-affinity A6C134–Tax interface. **(A)** Crystal structure of the A6C134–Tax complex (PDB: 4FTV). Left: ribbon diagram of the TCR–pMHC complex with CDR loops labeled. TCRα and TCRβ CDR loops are shown in pink and orange, respectively. Right: close-up of CDR3β residues L98, M99, S100, and A101 (blue) contacting TAX peptide residues V7 and Y8 (red). **(B)** Representative flow cytometry plots showing ZsGreen1 reporter activation in uninduced library cells (baseline), at the first round of droplet encapsulation and activation (R1), and at the second round of droplet encapsulation and activation (R2). **(C)** Enrichment score plotted against read count ranking for the top 100,000 TCR–pMHC pairs by read count. The wild-type LMSA–VY pair (red) is highlighted. **(D)** Bar charts showing read counts distributions of the top 100 TCR clonotypes after R1 enrichment. Clonotypes selected for experimental validation are indicated by red bar. **(E)** Pie charts showing the fraction of LMSA CDR3β clonotypes in the initial library, after first round of droplet-based enrichment (R1), and after second round of droplet-based enrichment (R2). **(F)** Scatter plots showing enrichment score (y-axis) plotted against read count ranking (x-axis) for peptide partners of TCR-LMSA (left) and TCR partners of peptide LLFGYPVYV (right). Wildtype pairs (VY and LMSA, respectively) highlighted in red; other experimentally tested variants shown in blue. **(G)** Bar charts showing %CD69 upregulation of the wild-type A6C134 TCR-LMSA against peptide variants spanning a range of read-count ranks (left), and of TCR variants spanning a range of ranks against the wild-type Tax peptide (right). Peptides were expressed as single-chain trimers on T2 cells. Data are presented as mean ± SD; n = 3 independent biological replicates. **(H)** Heatmaps of %CD69+ ratio (left) and counts number (right) for a matrix of five tested TCR variants against 11 peptide variants. Bold residues indicate the variant positions on peptide sequence. **(I)** Scatter plot of TCR cross-reactivity for all enriched TCR sequences. Each dot represents a unique TCR clonotype, plotted by the number of MHC variants recognized (x-axis, log2 scale) against fractional specificity (y-axis), defined as the fraction of read counts contributed by the single most abundant peptide partner. **(J)** Mutual information matrix quantifying statistical coupling between four CDR3β positions (rows, TCR positions 98–101) and two peptide contact positions (columns, peptide positions 7–8). **(K)** Atomic contacts between A6C134 CDR3β and the Tax peptide. CDR3β residues (L98, M99, S100, A101) contact Tax residues V7 and Y8, with dashed lines marking contacts within 4 Å (red, side-chain interaction; gray, main-chain interaction). Bottom: expanded views of the L98–Y8 and S100/M99/A101/L98–V7 contacts. **(L)** Sequence logos of CDR3β positions (T1–T4) stratified by peptide contact identity (P1, P2). Top: both peptide positions randomized (XX). Middle: the first position fixed as V (VX). Bottom: both positions fixed as V and Y (VY, wild-type Tax). Progressive restriction of peptide identity reveals increasing convergence of the CDR3β sequence landscape.

Enrichment analysis after both rounds revealed that the wild-type LMSA–VY pair ranked far outside the top-enriched combinations, a pattern distinct from the DMF4–MART-1 screen, where the wild-type pair dominated early rounds (**Figure 6C**). This observation is consistent with the broader functional sequence space accessible at a high-affinity interface, where many diverse sequences can satisfy the energetic threshold for TCR activation. When clonotypes were ranked individually, the wild-type LMSA CDR3β clonotype ranked first among enriched TCRs (**Figure 6D**), with modest enrichment rising from 0.04% in the starting library to 0.2% after R1 and 0.3% after R2 (**Figure 6E**).

Functional validation was then applied to selected pairs spanning a range of read counts with individual TCRs expressed in Jurkat cells and tested against their target peptides displayed as single-chain trimers on T2 cells, reading out activation by endogenous CD69 upregulation. First, activation was assessed by testing peptide variants of varying rank against the wild-type A6C134 TCR-LMSA (**Figures 6F, 6G**). Among the peptides tested, MG, EY, IS, and DY each induced strong CD69 upregulation comparable to the wild-type VY, whereas DG induced weak activation (∼6%), consistent with the low rank of VY among all LMSA- paired peptides in the selected pool (**Figure 6G**). In parallel, TCR variants were ranked against the wild-type VY peptide: LVSV, VFSS, and LVGN, all drawn from the top 100 TCR clonotypes, each showed wild-type VY peptide as their top binder and induced over 90% CD69 upregulation against the wild-type Tax peptide, while the lower-count FSIV also preferred VY and with weaker (∼50%) activation (**Figures 6D, 6F, 6G**).

Specificity was next mapped systematically by testing the five TCR variants against 11 peptide variants, enabling direct comparison of functional activation with library enrichment (**Figure 6H**, **S7C-I**). Wild-type LMSA was activated by 9 of 11 tested peptide variants, reflecting the broad cross-reactivity expected of this high-affinity wild-type, and matching its behavior in the paired sequencing analysis (**Figure 6H**). By contrast, LVSV, VFSS, LVGN, and FSIV showed narrower reactivity profiles, responding primarily to VY and EY. The activation was broadly concordant with the counts, with a few exceptions: DY and MS accumulated high read counts for VFSS yet induced only moderate CD69 activation, while FI and WL induced moderate activation of 30-40% across multiple TCRs despite low read counts in the count matrix (**Figure 6H**, **S7D-I**). Across all tested pairs, CD69 activation correlated with read count (Spearman ρ=0.71, p=1.43e-09) and with enrichment score (ρ=0.435, p=1.61e-09) (**Figure S7J**), confirming broad consistency between pair enrichment and experimental activation.

Across the full enriched pool, the extent of TCR cross-reactivity varied substantially. Fractional specificity, which reflects how selectively a TCR engages its preferred peptide partner relative to all others, was calculated for each enriched clonotype. Most TCR clonotypes recognized only a small number of peptide variants at fractional specificity above 0.3, whereas a minority, including the wild-type LMSA TCR, showed broad cross-reactivity (**Figure 6I**). The enriched repertoire, therefore, skews toward more constrained peptide specificity than the wild-type A6C134 TCR. Beyond individual specificity, the coevolution of the two chains was examined through a mutual information matrix computed between CDR3β positions 98–101 and peptide contact positions 7 and 8 (**Figure 6J**). Among all coupled position pairs, CDR3β residues L98 and M99 covaried most strongly with both peptide positions, whereas A101 covaried least with Tyr8. This pattern follows the physical interface: in the A6C134–Tax structure (PDB: 4FTV), L98 makes direct van der Waals contacts with Tyr8 of the Tax peptide, while A101 is positioned farthest from Tyr8 (**Figure 6K**). The mutual information coupling therefore tracks the physical proximity of these residues, indicating that the statistical interdependence captured by the library screen mirrors the underlying atomic organization at the TCR–pMHC interface.

The same coupling was visualized directly by stratifying CDR3β sequence logos according to peptide contact identity, which revealed how peptide sequence progressively constrains the compatible TCR sequence space. When both peptide contact positions were unrestricted, all four CDR3β positions showed broad amino acid diversity. Fixing the first peptide position to V narrowed the distribution, enriching the hydrophobic residues F and L at CDR3β position 1. Constraining both positions to VY converged the sequence space further, with position 1 strengthening its preference for F and L and position 2 showing enrichment for M and L (**Figure 6L**). Consistent with the crystal structure, the hydrophobic Tyr8 is packed against L98 and M99 and engaged directly by L98, accounting for the selection of hydrophobic residues at both positions to preserve this packing (**Figure 6K**). Together, these results demonstrate that individual peptide residues exert hierarchical, position-specific selective pressure on compatible CDR3β sequences, and that the LINC-seq platform captures these constraints at single-residue resolution across millions of paired variants (**Figure 6L**).

## Discussion

A longstanding bottleneck in functional TCR-peptide-MHC mapping has been the technical challenge of sequencing both partners from cell pairs engaged in productive signaling in a native context. LINC-seq addresses this by replacing unstable T cell/APC cell–cell doublets with genetically identical daughter cells derived from clonal expansion of a single founder within isolated semi-permeable capsules. This capability enables library-on-library co-variation across both interacting partners simultaneously. With this approach, any cell-surface receptor–ligand pair whose engagement drives transcriptional output is now accessible to function-based, library-scale screening using the same workflow. Cell-surface interactions are governed by epistasis between sequence variation in the receptor and in the ligand, and the rules of that epistasis can now be read by co-varying both.

TCR–peptide-MHC recognition is a stringent test case of any cell–cell receptor–ligand platform. TCR– pMHC output is a product of epistasis: small changes in either the CDR3 loop or in peptide anchor residues disrupt or rescue specificity in ways that cannot be predicted from either partner alone. Furthermore, the de-orphanization gap (10^15^–10^18^ TCRs in human repertoires, with cognate antigens known for only a few hundred) is a practical ceiling on TCR-based therapeutics^43^. A platform that captures TCR–pMHC epistasis solves a representative high-bar case, and the same workflow can be applied unchanged to less diverse receptor–ligand families (e.g. IgSF, GPCR, etc.).

Three experiments establish that the platform performs as designed in different regimes. First, cognate-pair recovery from a 3×3 pooled library at >95% fidelity confirms that trans engagement between clonal daughters in capsules faithfully recapitulates known TCR–pMHC specificity, and that paired sequencing from a single sort correctly recovers both partners. Second, simultaneous library-on-library deep mutational scans of three TCR–pMHC interfaces, co-varying up to ∼10^7^ joint variants per screen, rediscovered each wild-type clonotype among the top-ranked sequences and mapped the co-evolutionary fitness landscape across both partners in a single experiment, which one-sided mutagenesis cannot achieve. Third, screening interfaces of low, moderate, and high affinity (DMF4–MART-1, TCR55–Pep20, and A6C134–Tax) revealed that the breadth of the functional landscape scales with affinity: selection converged sharply onto the wild-type sequence at the low-affinity interface, tolerated a broader set of variants at moderate affinity, and produced an expansive landscape at high affinity in which the wild-type pair was only one of many functional pairs. These results show that a low-affinity interface lies near the activation threshold and tolerates little substitution, whereas a high-affinity interface retains surplus binding energy that can accommodate many suboptimal but still-sufficient peptide-contact sequences.

Beyond validating the platform, the co-evolution of these TCR-pMHC interfaces, revealed by two-sided mutagenesis, is informative. At the high-affinity A6C134–Tax interface, broad cross-reactivity was confined to a minority of enriched clonotypes that included the wild-type, and several of the narrower, more selective clonotypes were validated functionally. More broadly, each screen reports both which TCRs engage a given peptide and the cross-reactivity profile of a given TCR across the full panel of peptide variants, information directly useful for engineering target-specific TCRs for therapeutic use. The map also revealed structured interactions within the interface: the statistical coupling between receptor and peptide positions tracked the physical proximity of the contacting residues in the A6C134–Tax structure, linking the functional map directly to the atomic interface.

The most immediate, broader application in TCR biology is the de-orphanization of TCRs from tumor-infiltrating lymphocytes (TILs). Single-cell TCR sequencing has cataloged thousands of TIL clonotypes across human cancers^16–18,44^, but the antigens recognized by the overwhelming majority remain unknown and inferring them computationally has not been reliable^45,46^; pairing an unbiased peptide library against TIL TCR libraries in our platform reduces this to a single screening campaign. The same workflow extends to any cell-surface receptor–ligand family whose engagement drives a transcriptional output: immunoglobulin-superfamily inhibitory and co-stimulatory checkpoints, many of which remain functionally orphaned. This includes natural-killer-cell receptors, BCR–antigen pairs, and growth-factor or cytokine receptor families that converge on NFAT, NFκB, AP-1, or STAT transcription. We anticipate that the platform’s central design principle of converting cell–cell engagement into a single-cell-resolvable state will generalize widely across receptor–ligand biology.

We note several limitations that define the platform’s current operating envelope. The reporter is a Jurkat-derived T cell line, which does not fully recapitulate the activation thresholds and effector biology of primary CD8⁺ T cells; extension to primary cells will be needed to capture selection driven by metabolic state, exhaustion, or co-receptor engagement. The platform reads activation rather than killing, and follow-up cytotoxicity assays remain necessary for top candidates. The current 10⁷-variant ceiling is set by capsule throughput and FACS recovery rather than by a theoretical limit; further scaling is achievable with higher-throughput or parallel droplet generation and by isolating reporter cells without FACS. Finally, the current system uses a transcription-coupled reporter, and different reporter paradigms will be needed to extend the platform to the small subset of receptor–ligand pairs whose engagement does not converge on a transcriptional output.

## Materials and methods

### Cell Lines

Jurkat E6 TCR knockout cells were maintained in RPMI-1640 supplemented with 10% fetal bovine serum (FBS), 2 mM L-glutamine, 100 U/ml penicillin, and 100 μg/ml streptomycin at 37°C and 5% CO2. K562 cells and T2 cells were maintained under the same conditions. All cell lines were confirmed mycoplasma-negative prior to experimental use. No human subjects or clinical samples were used in this study.

### Semi-Permeable Capsule Fabrication

#### Capsule screening

Dextran-PEG-Mal droplets formed DTT-mediated core-shell structures in oil (**Figure S3B**) but were effectively irreversible under culture conditions, thereby limiting cell retrieval. Dextran- PEG-OPSS also formed capsules (**Figure S3B**), and the shell was removable by DTT-mediated reduction (**Figure S3C**). However, cells grown in Dextran-PEG-OPSS capsules decreased in viability over multi-day culture (**Figure S3D**), and encapsulated cells were frequently associated with or attached to the PEG-rich shell rather than undergoing uniform clonal expansion (**Figure S3E**). Dextran-PEGDA capsules also formed core-shell droplets with photo-initiator/UV-mediated crosslinking, but cells partitioned into the PEG-rich phase and were released outside the capsules after droplet breaking and transfer into culture media, indicating loss of cellular confinement (**Figure S3F**).

#### SPC generation

SPCs were generated using a jet-triggered, microfluidic device. The shell reagent was prepared by mixing 20% functionalized dextran solution with complete RPMI medium at a 1:1 ratio. The cell-containing core reagent contained 10% dextran-500K, 1.5% photoinitiator, and cells in complete medium. The cell concentration in the core reagent was adjusted at λ = 0.1 cells per 50 µm droplet to favor single-founder encapsulation.

For encapsulation, the shell reagent and core reagent were loaded into separate syringes and flowed into the jet-triggered microfluidic device at 800 µL/hr each. HFE-7500 oil containing 5% IK surfactant was introduced at 1500 µL/hr. Air was supplied to the air channel at 15 psi, generating periodic air bubbles that segmented the aqueous jet and triggered droplet break-off to produce monodisperse core-shell emulsions. The emulsion was collected for 1 h.

After collection, the emulsion was exposed to 405-nm wavelength light for 90 s to cure the shell. Capsules were immediately transferred from the oil to the aqueous phase by breaking the emulsion with 20% v/v 1H,1H,2H,2H-perfluoro-1-octanol (PFO) in HFE-7500, followed by centrifugation and washing with complete medium. Recovered SPCs were cultured in complete RPMI at 37°C and 5% CO_2_ for clonal expansion before Dox induction and downstream sorting. Viability of encapsulated cells was monitored by LIVE/DEAD Cell Imaging Kit.

#### TCR activation reporter Construct

Synthetic regulatory elements were designed by assembling varying copy numbers of NFAT-binding repeats (4*×*, 8*×*, 12*×*)^23^, NR4A motifs (12*×*)^24^, and other synthetic motifs^24^ are paired with one of three minimal promoters: the IL-2 minimal promoter (IL2miniP), the CMV minimal promoter (CMVminiP), or a synthetic minimal promoter (SMP)^24^. Each element was cloned upstream of the mCherry or ZsGreen1 fluorescence reporter gene in a lentiviral backbone.

#### TCR and pMHC Expression Constructs

The pLVX-TetOne-Puro-GFP plasmid was a gift from Jason Sheltzer (Addgene plasmid #171123). Peptide single-chain trimers (SCTs) were constructed in a single open reading frame with following orders: β2m leader sequence–peptide–GS linker–β2m–MHC α1α2α3. For the two-vector configuration, TCRα and TCRβ chains were cloned together with a P2A self-cleaving peptide sequence into the backbone of pLVX-TetOne-Puro-GFP lentiviral vector, under the control of either constant promoter or a doxycycline-inducible TRE3G promoter (Tet-On 3G system)^28^. Peptide single-chain trimers were cloned individually under TRE3G promoter. For the linked single-vector configuration, the pMHC SCT and TCRβ chain were co-expressed from a single TRE3G-driven vector via a P2A sequence, with TCRβ placed upstream of the pMHC SCT to achieve optimal surface expression. Human CD8α and CD8β subunits were co-expressed from a separate constitutive lentiviral vector (EF1α promoter).

#### Lentiviral Particle Production

Lentiviral particles were produced by co-transfecting HEK293T cells seeded at 1.5×10^6^ cells per well in a 6-well plate the day before transfection (70–90% confluency at time of transfection). For each well, a DNA master mix was prepared by diluting 2.5 μg total DNA in 125 μl OptiMEM (comprising 0.25 μg pMD2.G, 0.75 μg psPAX2, and 1.5 μg transfer plasmid) and adding 5 μl P3000 reagent. Separately, 5 μl Lipofectamine 3000 was diluted in 125 μl OptiMEM and incubated for 5 minutes at room temperature. The DNA and Lipofectamine mixtures were combined 1:1, incubated for 10–15 minutes at room temperature, and added to cells. After 6 hours, medium was replaced with fresh DMEM. Virus-containing supernatants were harvested at 48 and 72 hours post-transfection, filtered through 0.45 μm membranes, and used immediately or stored at −80°C.

#### Reporter cell line construction

Jurkat T cells were sequentially transduced with lentiviral particles to generate stable reporter lines. Target cells were counted and resuspended at 1×10^6^ cells in 500 μl cell culture medium, mixed with 1 ml viral supernatant, and transferred to a 24-well plate. Polybrene was added at a final concentration of 1 μg/ml, and cells were spun at 1500 g for 1.5 hours at 30°C. After spin infection, cells were incubated at 37°C and 5% CO_2_ for 4–6 hours, then transferred to fresh medium.

Cells were first transduced with the 8*×*NFAT-ZsGreen1-PGK-tNGFR reporter construct, which co-expresses a truncated nerve growth factor receptor (tNGFR) under the PGK promoter as a selection marker, stable cells were isolated by fluorescence-activated cell sorting for tNGFR surface expression. The hCD8 co-expression vector was then introduced, followed by sequential stable integration of the doxycycline-inducible TCRα construct carrying a Blasticidin resistance marker and the Dox-inducible TCRβ–pMHC SCT construct carrying a Puromycin resistance marker. Stable cells were selected with Puromycin (1 μg/ml) and Blasticidin (10 μg/ml) for 7 days to establish the reporter cell line.

#### Screening of Reporter Response Elements

Individual response element constructs driving mCherry expression were transduced into Jurkat T cells stably expressing the YLQ7, A6C134, or DMF5 TCR using the spin-infection protocol described above. Transduced cells were co-cultured with K562 antigen-presenting cells stably overexpressing the cognate peptide in SCT format at a 1:1 effector-to-stimulator ratio for 24 hours at 37°C. mCherry expression was quantified by flow cytometry, and fold-induction was calculated as the ratio of mCherry-positive cells in stimulated versus unstimulated conditions.

#### Small-scale Doxycycline induction of reporter cell

Reporter cells were seeded in 96-well U-bottom plates at 1×10^5^ cells per well in complete RPMI medium. Doxycycline was added at a final concentration of 100 ng/ml to induce co-expression of TCR and pMHC SCT, and cells were incubated at 37°C and 5% CO_2_ for 48 hours. Uninduced cells receiving no doxycycline were included in each experiment as a baseline control. ZsGreen1 reporter expression and surface marker staining were analyzed on a CytoFLEX flow cytometer (Beckman Coulter).

#### High-Throughput Encapsulation and Clonal Expansion

Reporter cells in mid-log phase were resuspended in core-phase polymer at 2×10^6^ cells/mL (Poisson loading ∼0.15 cells/capsule to ensure single-cell occupancy). After encapsulation, SPCs were transferred to T-175 flasks in complete RPMI-1640 and cultured at 37°C, 5% CO₂. Capsules were maintained in suspension culture. On day 2, 50% media exchange was performed by pelleting SPCs at 200 × g and resuspending in fresh media. Viability of encapsulated cells was monitored by calcein-AM / ethidium homodimer-1 staining on days 1 and 3.

#### Doxycycline Induction and FACS Sorting

On day 3 post-encapsulation, Dox was added to the culture media at a final concentration of 1 μg/mL. Induction proceeded for 48 hours at 37°C. SPCs were centrifuged at 300×g for 5 min, and washed with PBS. SPCs were then dissolved by addition of dextranase (2 µL of dextranase per 1 mL of PBS), and incubated for 5 min in room temperature. The resulting cell suspension was washed in PBS by centrifugation at 300×g for 5 min. ZsGreen1⁺ cells were sorted on a BD FACSAria III (530/30 filter).

#### Sanger sequencing validation of cognate pairs from bulk genomic DNA

For the 3x3 TCR–pMHC matrix validation, paired TCR–pMHC sequences were amplified from bulk genomic DNA of collected cells using a forward primer targeting the leader sequence region and a reverse primer targeting the TCR variable region. Amplicons of approximately 1.8 kb were generated using PrimeSTAR Max DNA Polymerase (Takara), cloned into the pLVX backbone, and transformed into competent bacteria. Individual colonies were picked, cultured overnight, and submitted for Sanger sequencing. Sequenced pairs were analyzed for cognate matching.

#### Single-cell RT-PCR validation of cognate pairs

Individual ZsGreen1-positive cells were sorted directly into 96-well PCR plates pre-loaded with 5 μl RT buffer mix and kept on ice. Single-cell reverse transcription was performed using the SuperScript IV First-Strand Synthesis System (Invitrogen) with random hexamers according to the manufacturer’s instructions, using an abbreviated thermocycler program (42°C 7.5 min, 23°C 7.5 min, 50°C 30 min, 94°C 3 min) to reduce processing time. cDNA was stored at −80°C or used immediately for downstream PCR amplification.

Paired TCR–pMHC sequences were amplified from single-cell cDNA by two-round nested PCR using gene-specific primers targeting the TCR variable region and the pMHC SCT sequence (primer sequences provided in Table S1). First-round PCR was performed using 2x HiFi mix with 1.5 μl cDNA as template for 45–50 cycles. Second-round PCR used 2 μl of 10-fold diluted first-round product as template for 40– 45 cycles. Amplicons were analyzed by agarose gel electrophoresis, purified, and submitted for Sanger sequencing.

#### Cloning of TCR-pMHC library

Combinatorial TCR–pMHC libraries were designed by simultaneously randomizing selected contact positions identified from published crystal structures. Primer oligos cover the target region with degenerate codons were synthesis by Integrated DNA Technologies (IDT). TCR–pMHC libraries were amplified and cloned into the pLVX lentiviral backbone using HiFi Assembly (NEB). Briefly, 110 fmol dsDNA insert and 20 fmol linearized backbone vector were combined with 10 μl HiFi Assembly Master Mix and incubated at 50°C for 1 hour (eight parallel reactions). Assembly products were pooled, purified using a PCR cleanup kit (Qiagen), and eluted in 30 μl water.

For bacterial transformation, MegaX DH10B T1R electrocompetent cells (Thermo Fisher Scientific) were thawed on ice. Purified assembly product (50 ng) was mixed with per 25 μl competent cells, transferred to a chilled 0.1 cm electroporation cuvette, and electroporated at 1.8 kV using a Bio-Rad Micropulser. Cells were immediately recovered in 1 ml SOC medium and incubated at 37°C with shaking at 225 rpm for 1 hour. Serial dilutions were plated on ampicillin LB agar plates and incubated overnight at 37°C to determine colony-forming units and verify library coverage. The remaining culture was transferred to 500 ml ampicillin 2×TY medium and grown overnight at 30°C with shaking at 225 rpm for 18–20 hours, after which plasmid DNA was extracted by Xtra Midi prep (MACHEREY-NAGEL).

#### TCR-pMHC library cell preparation

Reporter cells stably expressing the 8×NFAT-ZsGreen1 reporter and hCD8 were first transduced with the doxycycline-inducible TCRα construct using the viral infection protocol described above and selected with Blasticidin (10 μg/ml) for 7 days. The paired TCR–pMHC library lentiviral vector was subsequently introduced at low MOI (< 0.3) to ensure single-copy integration per cell, maintaining a one-to-one correspondence between TCR and pMHC identity. Library cells were selected with Puromycin (1 μg/ml) for 7 days and expanded to the required cell numbers prior to screening. A minimum of 300 million cells were used for library transduction to ensure coverage of library diversity.

#### TCR-pMHC library cell treatment in bulk with doxycycline

Library cells were cultured in complete RPMI medium and induced with doxycycline (100 ng/ml) for 48 hours at 37°C and 5% CO_2_ to drive co-expression of TCR and pMHC SCT. A total of 200 million library cells were used for each screening to ensure sufficient representation of library diversity. Uninduced control cells were maintained in parallel under identical conditions without doxycycline to establish the baseline ZsGreen1 negative population.

#### MACS enrichment for TCR-pMHC library cells

Following doxycycline induction, dead cells were removed using the Dead Cell Removal Kit (Miltenyi Biotec) according to the manufacturer’s instructions. CD69-positive activated cells were then enriched by magnetic-activated cell sorting using the CD69 Microbeads Kit II (Miltenyi Biotec). Briefly, cells were pelleted at 300×g for 10 minutes, resuspended in MACS buffer (PBS pH 7.2, 0.5% BSA, 2 mM EDTA) at 40 μl per 10^7^ cells, and incubated with CD69-Biotin for 15 minutes at 2–8°C. Anti-Biotin Microbeads were added and incubated for a further 15 minutes at 2–8°C. Cells were washed, resuspended in 500 μl MACS buffer per 10^8^ cells, and passed through an LS column on a MACS separator. The column was washed three times with 3 ml MACS buffer, then removed from the magnetic field and CD69 positive cells were eluted with 5 ml MACS buffer. Enriched cells were subsequently sorted for ZsGreen1 expression by FACS to obtain the final activated population for downstream sequencing and re-encapsulation.

#### Amplicon Preparation and Next-Generation Sequencing

Genomic DNA was extracted from the starting library and sorted cell populations using a custom lysis protocol. For larger cell numbers, cells were lysed overnight at 56°C in gDNA lysis buffer (200 mM NaCl, 50 mM Tris-HCl pH 7.5, 5 mM EDTA, 0.2% SDS), supplemented with Proteinase K (0.4 mg/ml final concentration). Genomic DNA was ethanol-precipitated, washed twice with 75% ethanol, air-dried, and dissolved in TE buffer by incubation at 56°C overnight. For smaller cell numbers, genomic DNA was extracted using the QIAamp DNA Mini Kit (Qiagen).

Paired TCR–pMHC amplicons were generated by two-round PCR using PrimeSTAR Max DNA Polymerase (Takara). Products were resolved by agarose gel electrophoresis, and the target band was excised and gel purified. First-round primers contained gene-specific sequences targeting the TCR–pMHC locus flanked by partial Illumina adapter sequences and sample-specific barcodes (primer sequences provided in Table S1). Second-round PCR was performed using 50–100 ng of purified first-round product as template for 10–12 cycles under the same thermocycler conditions, using Illumina P5 and indexed P7 primers to complete the full sequencing adapter sequences. Products were again gel-purified and quantified by Qubit fluorometry (Thermo Fisher) prior to sequencing.

For MiSeq sequencing, purified amplicons were sequenced on an Illumina MiSeq platform using a V3 reagent cartridge (paired-end, 250 bp) with 10% PhiX spike-in, following the standard Illumina library preparation protocol. For NovaSeq sequencing, amplicons were sequenced on an Illumina NovaSeq 6000 platform with paired-end 150 bp reads (Signios Biosciences).

#### NGS sequence analysis

Raw paired-end reads were processed using a custom bioinformatics pipeline. Adapter sequences were trimmed and read pairs lacking the expected adapter were discarded using cutadapt (v5.2). Reads were quality-filtered using fastp^47^ with a minimum Phred quality score of 20 and a maximum low-quality base fraction of 20% per read. Following quality filtering, additional flanking sequences were removed, and the CDR3β-encoding region was extracted from R1 reads and the peptide-encoding region from R2 reads using cutadapt. R2 reads were reverse-complemented using seqtk to obtain the sense-strand peptide sequence.

R1 and R2 reads were merged into a single sequence per read pair. Merged sequences were filtered by verifying alignment to expected CDR3 and peptide flanking sequences at defined positions, and read pairs failing this filter were discarded. The variable CDR3β and peptide positions within each passing read were translated into amino acid, and sequences containing premature stop codons were excluded. Unique TCR– pMHC amino acid pair combinations were enumerated and their read counts tabulated separately for CDR3β sequences, peptide sequences, and full paired combinations. Statistical enrichment of each TCR– pMHC pair was assessed using a one-sided hypergeometric test comparing the observed co-occurrence frequency of each pair against the marginal frequencies of the individual TCR and peptide sequences. Pairs with a p-value below 0.05 were considered significantly enriched.

#### Functional Validation Assay

Jurkat TCR knockout cells with an overexpressed 8*×*NFAT-ZsGreen1 reporter and hCD8 were used to validate the activation of the tested TCR variants. TCRs were packaged into lentivirus and used to infect Jurkat cells. T2 cells stably co-expressing BFP fluorescence protein and a single-chain trimer MHC with the target peptide sequence. Jurkat cells expressing individual TCR were mixed with T2 cells at a 1:1 effector-to-stimulator ratio. Co-cultures were incubated at 37°C and 5% CO_2_ for 12-16 hours. Cells were stained with anti-CD69-APC for 15 minutes and washed once then analyzed on a NovoCyte Quanteon flow cytometer (Agilent). Percent CD69-positive cells were calculated using FlowJo v10 with gating on BFP-negative (TCR-expressing Jurkat) cells. Data are presented as mean ± standard deviation of three biological replicates (n = 3).

#### Enrichment score calculation

The enrichment score was calculated using Enrich2 with the ratios scoring method^48^. The log ratio for pair i, equivalent to its log fold change, was calculated as the natural log ratio of read counts, each normalized to total library size, between the final and initial libraries.

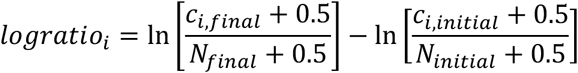

Where c_t_, t is the read count for pair *i* in library *t* (initial or final), and N_t_ is the total number of reads in library t. Normalizing each count by total library size accounts for differences in sequencing depth between the two libraries.

The variance of each score was estimated from Poisson counting statistics using the delta method, in which the variance of the log of a count is approximately the inverse of the count.

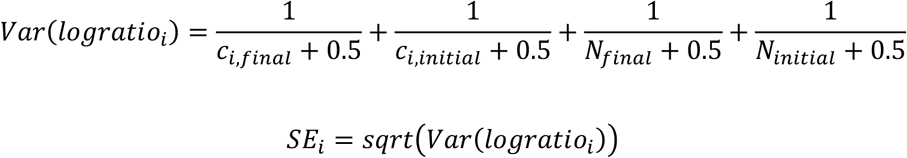

Log ratio enrichment score was divided by its standard error to produce a normalized enrichment score. Enrichment score values indicate pairs enriched in the final selected round relative to the initial library.

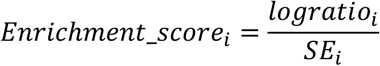

#### Sequence Logo Analysis

Sequence logos were generated using either the WebLogo (v3.7)^49^ or Python package logomaker^50^ with probability-based letter scaling and chemistry color scheme for amino acid sequences. A defined region was extracted from each translated FASTA file, and logos were computed separately for the starting library, round 1-sorted, and round 2-sorted populations to visualize progressive positional convergence across screening rounds.

#### Fractional Specificity Calculation

For each unique TCR clonotype recovered from the screen, fractional specificity was defined as the fraction of that TCR’s total read counts contributed by its single most abundant peptide partner, calculated as: fractional specificity = (read counts of most abundant peptide partner) / (total read counts across all peptide partners). Values approaching 1.0 indicate high peptide selectivity; values approaching 0 indicate broad cross-reactivity.

#### Mutual Information Matrix Computation

Mutual information (MI) between CDR3β positions and peptide contact positions was computed from the joint amino acid frequency distributions of all enriched TCR–pMHC pairs recovered from R2-sorted cells.

For each position pair (i, j), MI was calculated as

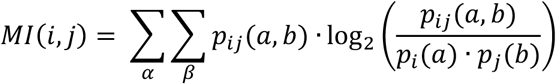

where *a* and *b* index the amino acids observed at positions *i* and *j*, *p_i_*(*a*) and *p*_j_(*b*) are the marginal amino acid frequencies at positions *i* and *j*, and *p_i_*_j_(*a*, *b*) is their joint frequency. MI was computed for all combinations of CDR3β positions 98–101 against peptide contact positions 7 and 8, yielding a 4×2 matrix. Statistical significance was assessed against a permutation null distribution generated by randomly shuffling TCR–pMHC pair assignments across 1,000 permutations.

#### Sequence similarity network analysis

For each sample, peptide and TCR sequences corresponding to the same pair were averaged by read count and concatenated into a single sequence representing the peptide and TCR pair. The 1000 most abundant pairs were selected for network analysis. Pairwise edit distances between all selected sequences were calculated using the Levenshtein distance, and an edge was drawn between two sequences when their edit distance was less than 2. Network layout was computed using the Kamada Kawai algorithm based on this full set of edges. Node size was scaled in proportion to read count, and the target pair was highlighted in a distinct color to track its position within the network. Network construction and visualization were performed in Python using the igraph and python-Levenshtein packages.

#### Quantification and statistical analysis

All statistical analyses were performed in Python (v3.10; SciPy, NumPy) unless otherwise noted, and flow cytometry data were analyzed in FlowJo (BD Biosciences) as described above. Detailed statistical parameters, exact value of n and significance values were provided in the corresponding figures and figure legends. Data are presented as mean ± SD, with n = 3 independent biological replicates. Correlation between functional activation (%CD69+) and either sequencing read count or normalized enrichment score across experimentally validated TCR–pMHC pairs was assessed by two-sided Spearman rank correlation (read count: ρ = 0.71, p = 1.43×10^-9^; enrichment score: ρ = 0.435, p = 1.61×10^-9^)

#### Materials availability

There are restrictions to the availability of the functionalized dextran because of the lack of an external centralized repository for its distribution and our need to maintain the stock. We are glad to share the functionalized dextran with reasonable compensation by requestor for its processing and shipping. Plasmids generated in this study will be provided by the lead contact upon completion of a Materials Transfer Agreement.

#### Data and code availability

Sequencing data have been deposited at NCBI Sequence Read Archive (SRA) and will be publicly available as of the date of publication. The BioProject accession number is PRJNA1502447. All original code will be deposited at GitHub and will be publicly available as of the date of publication. Any additional information required to reanalyze the data reported in this paper is available from the lead contact upon request.

## Acknowledgements

We acknowledge Mark M. Davis for sharing of TCR knockout Jurkat cell line. KCG is an Investigator of the Howard Hughes Medical Institute, and also funded by Parker Institute for Cancer Immunotherapy,

Ludwig Cancer Institute, Yosemite Innovation Fund, and NIH R01GM150125. ICC is an Arc Innovation Investigator. ICC and SWS were supported by NIH (DP2AI154435).

## Author contributions

Conceptualization, L.D.L., S.W.S., I.C.C., and K.C.G.; Methodology, L.D.L., S.W.S., K.J., J.P.; Investigation, L.D.L., S.W.S., K.J., C.W., J.P., R.M., and X.X.; Writing – original draft, L.D.L., S.W.S., I.C.C., and K.C.G.; Writing – review & editing, all authors; Funding acquisition, I.C.C. and K.C.G.; Resources, I.C.C. and K.C.G.; Supervision, I.C.C. and K.C.G. All authors reviewed and approved the final manuscript.

## Declaration of generative AI and AI-assisted technologies in the manuscript preparation process

During the preparation of this work, the authors used Claude and ChatGPT in order to improve the grammar, language, and readability of the manuscript. After using the tools, the authors reviewed and edited the content as needed and take full responsibility for the content of the published article.

---

**Figure S1.**
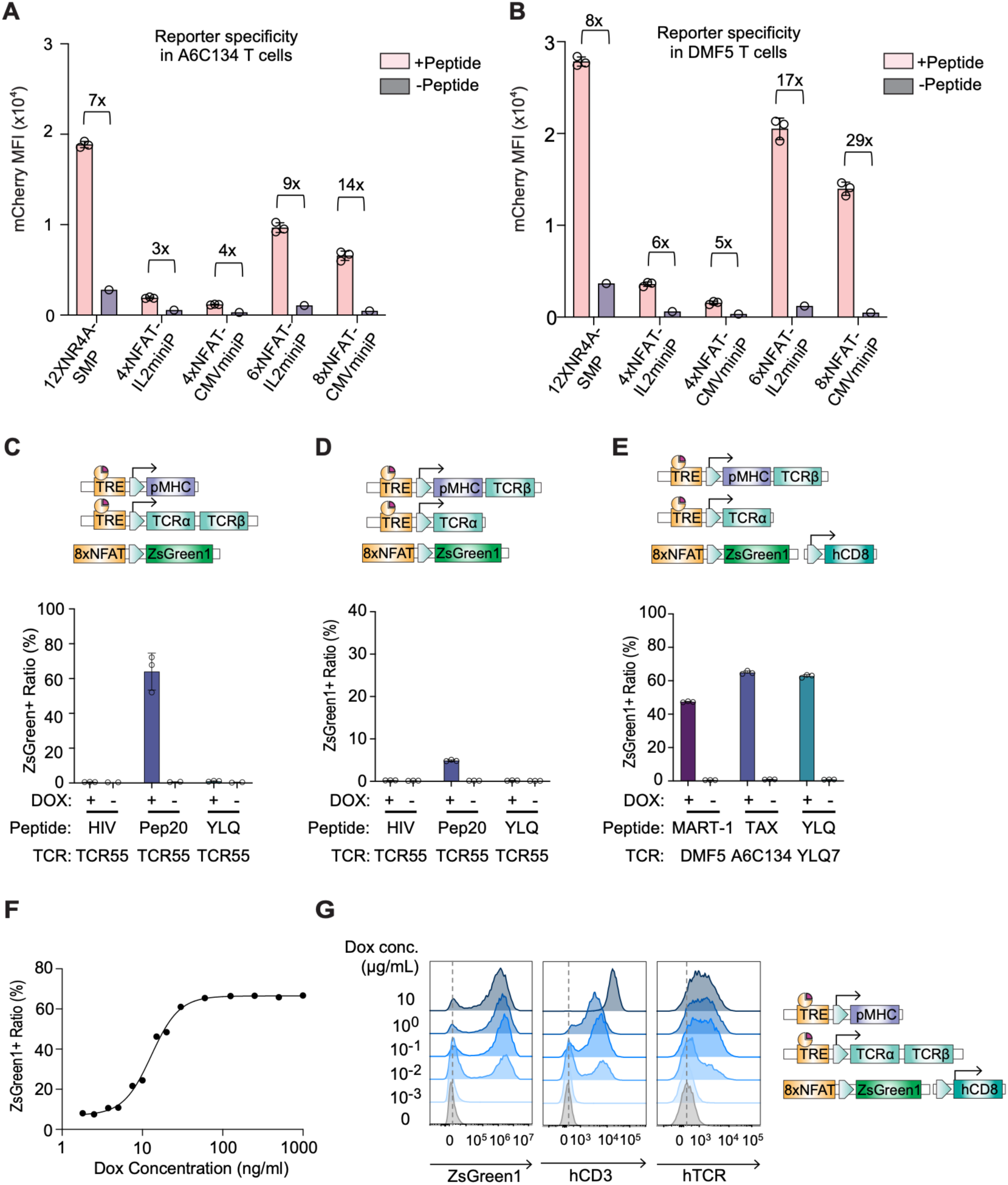
Additional characterization of the LINC-seq reporter system. **(A–B)** Response elements driving mCherry expression were screened in Jurkat T cells expressing A6C134 **(A)** and DMF5 **(B)** TCRs, co-cultured with antigen-presenting cells (APCs) presenting the Tax or MART-1 peptide as a single-chain trimer (SCT), respectively. mCherry MFI is shown with (+Peptide) and without (–Peptide) cognate peptide; fold induction over the –Peptide baseline is indicated above selected bars. **(C–D)** 8xNFAT-ZsGreen1 reporter cells without hCD8 co-expression were tested against the moderate agonist (Pep20), weak agonist (HIV), and non-cognate (YLQ) peptides (all paired with TCR55), with TCRα and TCRβ co-encoded on a single construct **(C)** or TCRβ fused to the pMHC and TCRα on a separate construct **(D)**. Schematics indicate construct architecture. **(E)** Reporter cells co-expressing hCD8 and a single-construct pMHC-TCRβ were validated across additional TCR–peptide pairs: DMF5/MART-1, A6C134/Tax, YLQ7/YLQ. **(F)** 8xNFAT-ZsGreen1 reporter cells with constitutive TCR expression and dox-inducible pMHC were treated with a 10-fold Dox titration. ZsGreen1-positive ratio was fitted to a sigmoidal dose-response curve. **(G)** Flow cytometry histograms of ZsGreen1, hCD3, and hTCRα/β expression in reporter cells carrying pMHC and TCR under separate Dox-inducible promoters, across a Dox titration (0–10 µg/mL), showing dose-dependent induction that saturates at 100 ng/mL. Data in (A)-(E) are presented as mean ± SD; n = 3 independent biological replicates.

**Figure S2.**
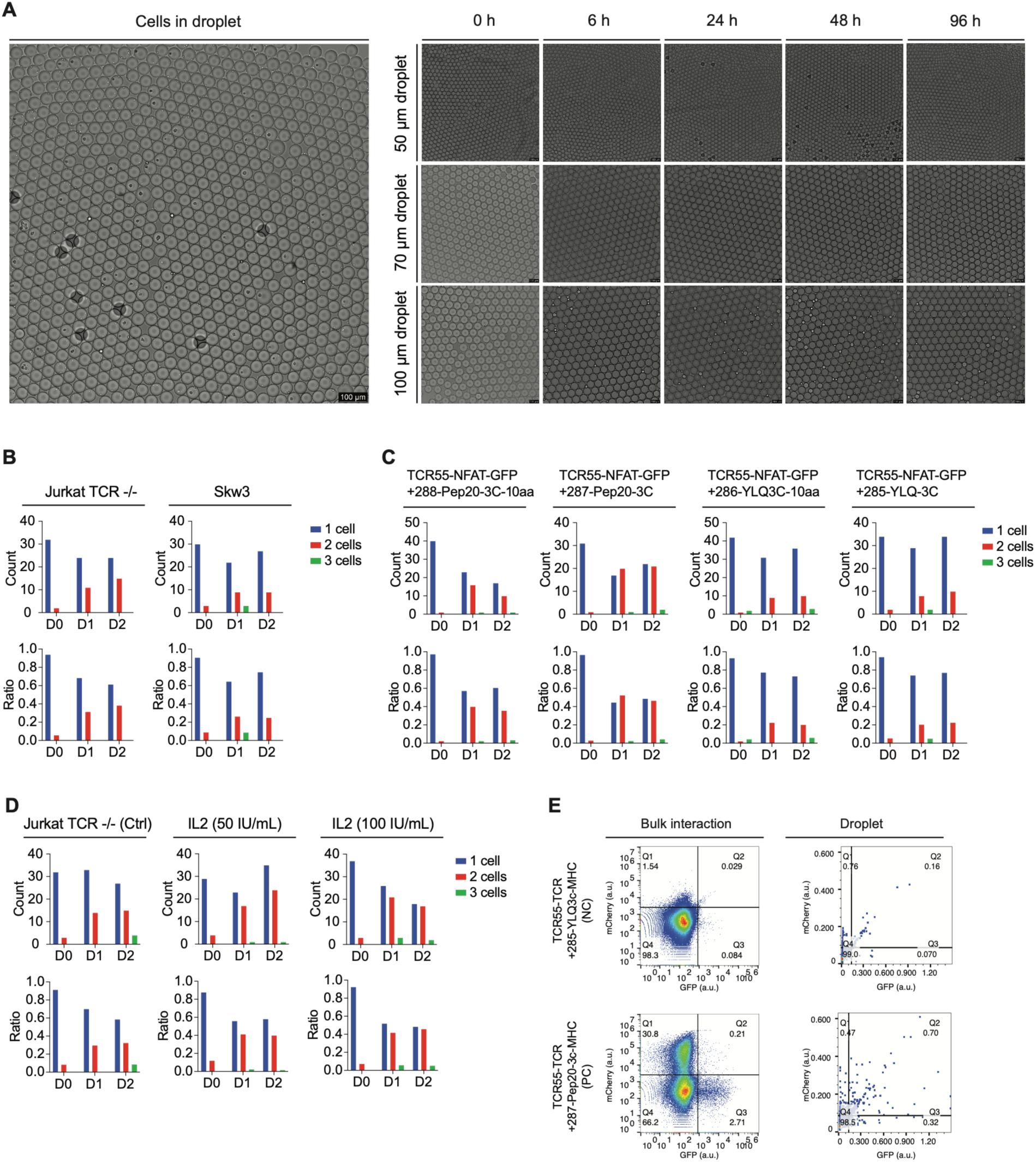
Cell culture in water-in-oil droplet compartmentalization. **(A)** Brightfield images of cells encapsulated in water-in-oil droplets. Left, representative image of cell-containing droplets. Right, representative droplets of nominal diameters of 50, 70, and 100 μm imaged over 0-96 h to assess droplet stability during extended culture. **(B)** Quantification of Jurkat TCR^−/−^ and SKW3 cell occupancy in droplets over three days. Bar plots show the number of droplets containing 1, 2, or 3 cells at day 0, day 1, and day 2; corresponding ratio plots show the fractional distribution of each occupancy class. **(C)** Droplet culture of TCR55-NFAT-GFP reporter cells carrying indicated pMHC constructs. Droplet occupancy was quantified over day 0-2 for cells expressing cognate Pep20 or non-cognate YLQ constructs of different linker/construct configurations. Bar plots show counts of droplets containing 1, 2, or 3 cells; ratio plots show normalized occupancy distributions. **(D)** Effect of IL-2 supplementation on droplet-contained Jurkat TCR^−/−^ cells. Cells were cultured without IL-2 or with 50 or 100 IU/mL IL-2, and droplet occupancy was quantified over day 0-2. IL-2 supplementation did not restore robust clonal expansion within droplets. **(E)** Comparison of NFAT-GFP reporter activation in bulk culture and droplet culture. Bulk interaction assays produced detectable NFAT-GFP (TCR-encoded cells) and NFAT-mCherry (pMHC-encoded cells) activation following Dox induction, whereas droplet-confined cells showed limited reporter activation, consistent with poor long-term culture performance in sealed droplets.

**Figure S3.**
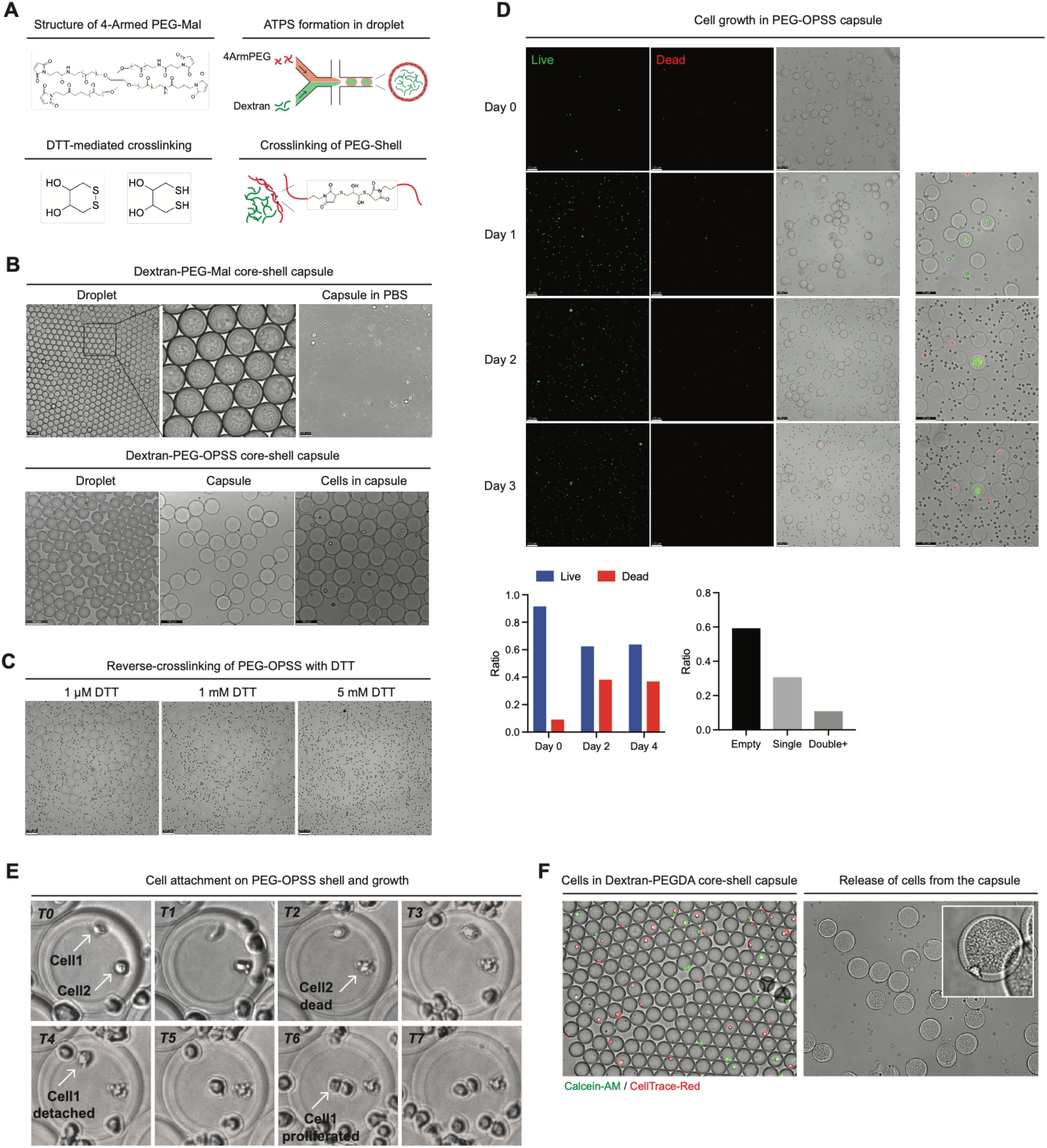
Evaluation of PEG-Dextran ATPS-based core–shell capsule chemistries. **(A)** Schematic of PEG-based core–shell capsule formation by aqueous two-phase separation (ATPS). Four-arm PEG-maleimide (4ArmPEG-Mal) and dextran phase-separate within droplets to generate a dextran-rich core and PEG-rich shell. The PEG shell can be crosslinked through thiol-maleimide chemistry, and disulfide-containing PEG crosslinkers enable DTT-mediated reverse crosslinking. **(B)** Brightfield images of dextran–PEG-Mal and dextran–PEG-OPSS core–shell capsule formation. **(C)** DTT-mediated dissolution of PEG-OPSS capsules. Jurkat cells were encapsulated in PEG-OPSS capsules, and additional free Jurkat cells were added to the surrounding culture medium to support cell viability and growth. Capsules were treated with 1 μM, 1 mM, or 5 mM DTT to test reduction-triggered shell disruption. Higher DTT concentrations promoted capsule destabilization and release of encapsulated contents, confirming partial reversibility of the disulfide-crosslinked PEG-OPSS shell. **(D)** Cell viability and growth in PEG-OPSS capsules over three days. Live/dead fluorescence images and merged brightfield images show encapsulated cells from day 0 to day 3. Quantification of live and dead cell fractions indicates reduced viability over extended culture. Occupancy analysis shows the distribution of empty, single-cell, and doublet-containing capsules. **(E)** Time-lapse brightfield imaging of cell behavior within PEG-OPSS capsules. Encapsulated cells frequently attached to the PEG-OPSS shell, and individual cells showed divergent outcomes, including detachment, death, and limited proliferation. These observations indicate that PEG-OPSS capsules were not optimal for uniform clonal expansion of Jurkat. **(F)** Dextran–PEGDA core–shell capsules failed to retain Jurkat cells within the dextran-rich core after transfer to aqueous culture media. Left, cell-staining image shows that cells were initially co-encapsulated within droplets containing Dextran–PEGDA core–shell structures. However, many cells partitioned into or became associated with the PEG-rich shell phase rather than remaining confined in the dextran-rich core. Right, after droplet breaking and transfer of capsules into culture media, cells were released outside the capsule structures, indicating poor cellular retention.

**Figure S4.**
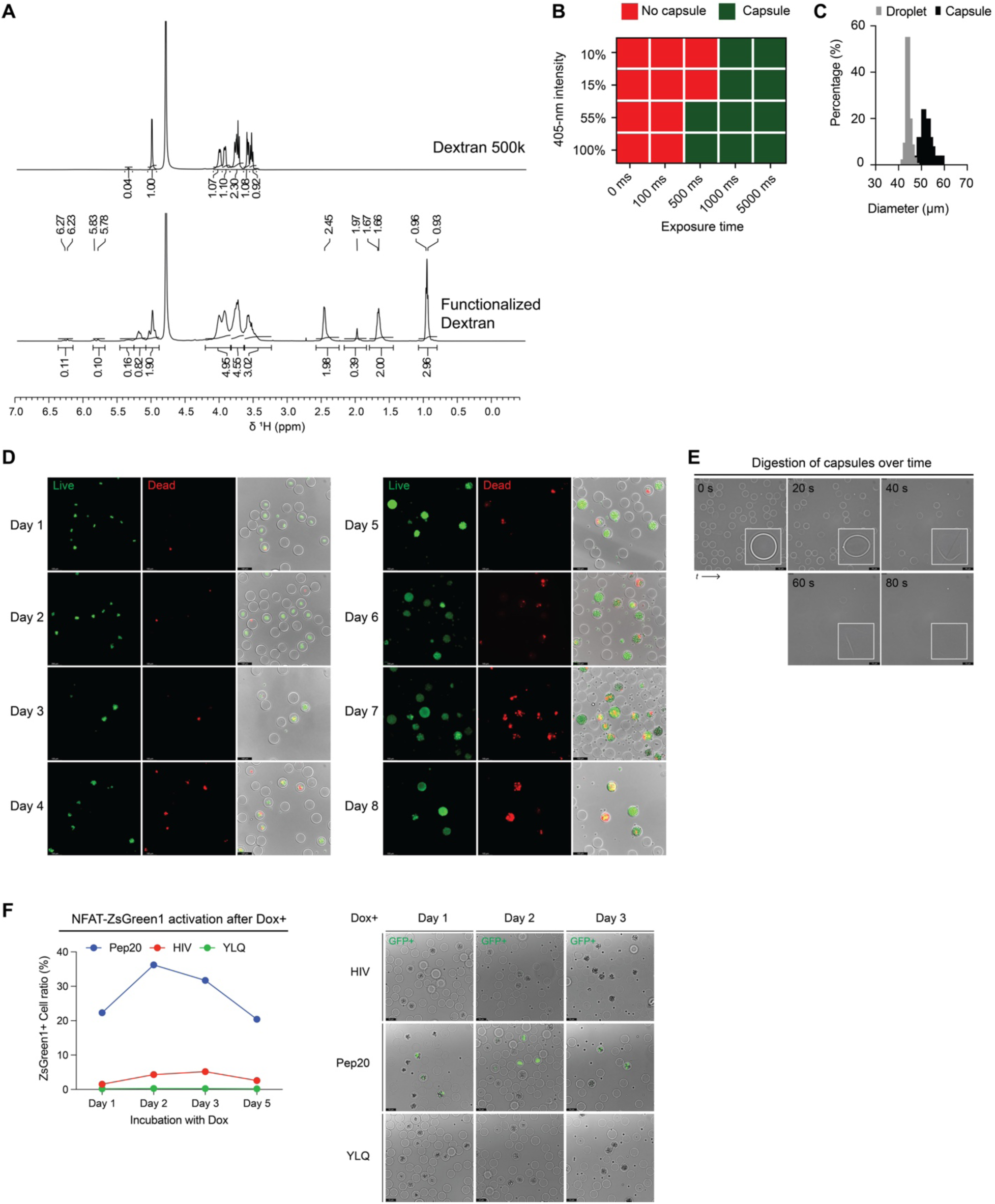
Synthesis, optimization, and functional validation of dextran-based semi-permeable capsules. (A) ^1^H NMR spectra of unmodified dextran 500K and functionalized dextran in D_2_O. Compared with dextran 500K, functionalized dextran showed diagnostic methacrylate vinyl resonances at δ 6.27/6.23 and 5.83/5.78 ppm, a methacrylate methyl resonance at δ 1.97 ppm, and butyrate resonances at δ 2.45, 1.67/1.66, and 0.96/0.93 ppm, confirming co-functionalization of dextran with methacrylate and butyrate groups. (B) 405-nm exposure intensity and exposure-time matrix for shell curing. Red indicates conditions that failed to produce stable capsules after transfer to aqueous medium, and green indicates conditions that produced stable capsules. Final collected emulsions were cured by 405-nm illumination for 90 s. (C) Size distributions of droplets before shell curing and recovered capsules after transfer into aqueous medium. (D) Live/dead staining of Jurkat cells cultured in SPCs over eight days. Live cells are shown in green and dead cells in red. Merged brightfield/fluorescence images show maintenance of capsule-confined colonies during extended culture. (E) Time-lapse brightfield images of capsule digestion. Insets show representative single capsules during shell disruption and loss of capsule structure over time. (F) NFAT-ZsGreen1 reporter activation in SPCs after Dox induction. Reporter cells encoding moderate cognate TCR55–Pep20, weak cognate TCR55–HIV, or non-cognate TCR55–YLQ pairs were encapsulated, expanded, induced with Dox, and imaged over time. Jurkat TCR^−/−^ cells were added to the surrounding culture medium to support cell viability and growth. After 3 days of culture, cells outside the capsules were removed using a 40-µm cell strainer. Quantification shows selective ZsGreen1 activation in the moderate cognate condition, with reduced activation in the weak cognate condition and minimal activation in the non-cognate control.

**Figure S5.**
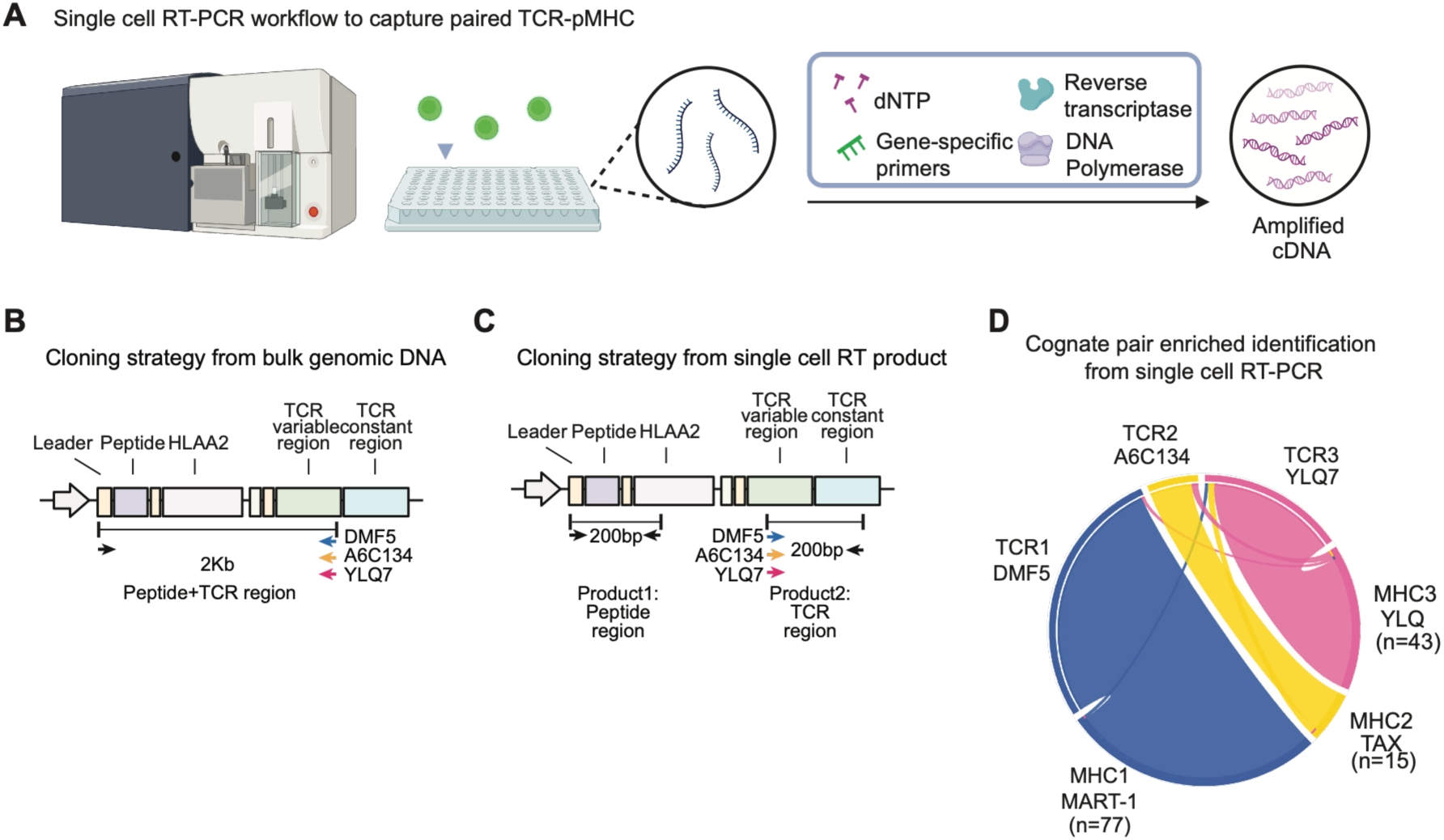
Validation of paired TCR–pMHC identification by single-cell RT-PCR. **(A)** Schematic of the single-cell RT-PCR workflow for recovering paired TCR–pMHC sequences. Single cells are isolated, and their mRNA is reverse-transcribed and amplified with gene-specific primers, reverse transcriptase, and DNA polymerase to generate paired cDNA. **(B–C)** Cloning strategies for paired TCR–pMHC amplification from bulk genomic DNA **(B)** or single-cell RT products **(C)**. From bulk gDNA, a single ∼2-kb amplicon spans the peptide and TCR regions; from single-cell RT, two ∼200-bp products are generated separately for the peptide and TCR regions. Schematics indicate primer positions and expected product sizes. **(D)** Chord diagram of cognate TCR–pMHC pairs recovered from ZsGreen1-positive sorted cells by single-cell RT-PCR. n indicates the number of pairs recovered per cognate pMHC.

**Figure S6.**
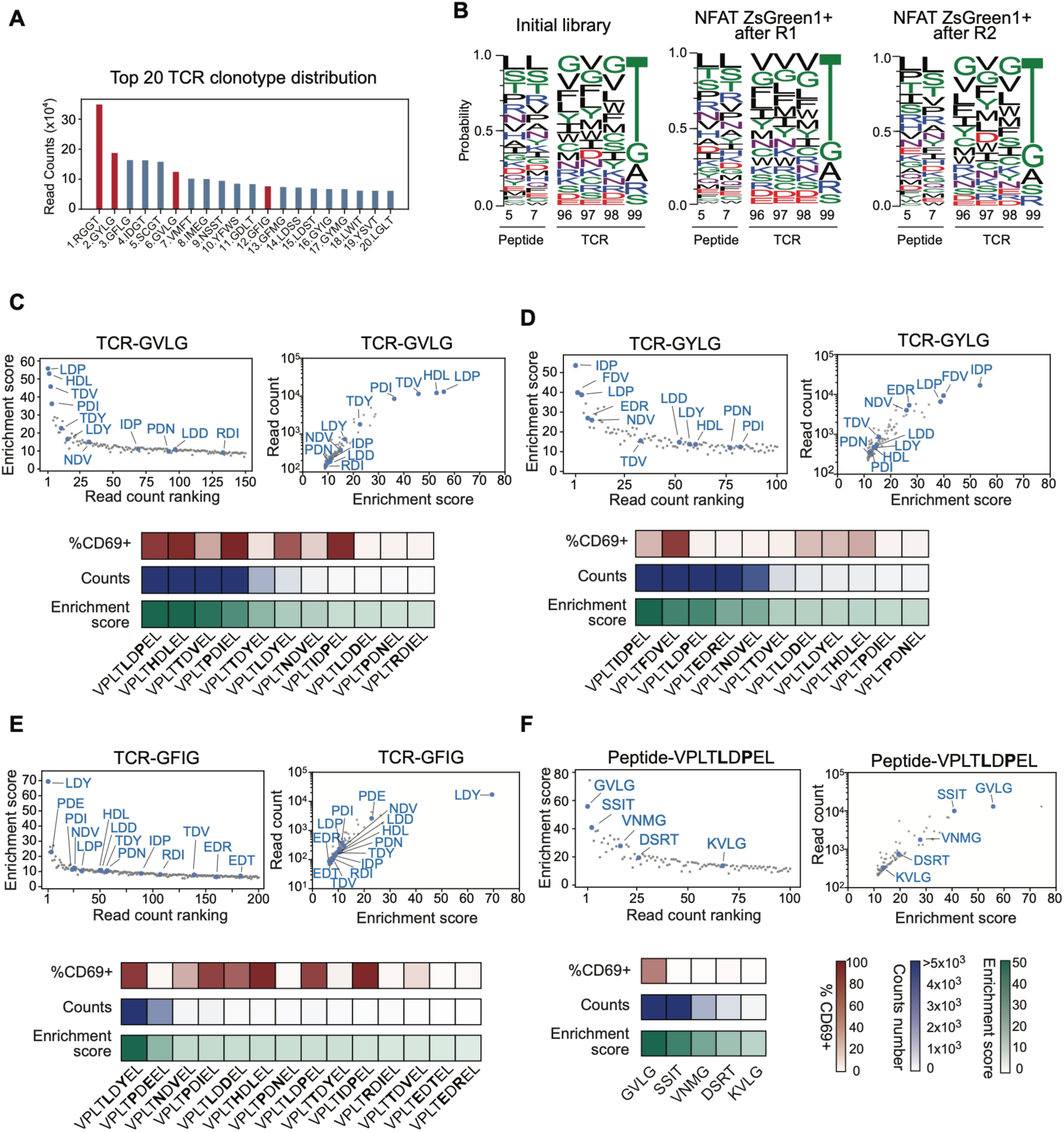
LINC-seq enrichment and validation of the Pep20 peptide–CDR3β library. **(A)** Read-count distribution of the top 20 TCR clonotypes after enrichment. Clonotypes selected for experimental validation (RGGT, GYLG, GVLG, and GFIG) are highlighted in red. **(B)** Sequence logos of the mutagenized peptide positions (5 and 7) and CDR3β positions (96–99) in the initial library, after bulk sorting (R1), and after SPC-based sorting (R2). **(C–F)** Characterization of top-ranked TCR–pMHC pairs for TCR-GVLG (C), TCR-GYLG (D), TCR- GFIG (E), and the TCR partners of peptide VPLTLDPEL (F). Left scatter plots show enrichment score (y-axis) plotted against read count ranking (x-axis). Right scatter plots show read count (y-axis) plotted against enrichment score (x-axis). Heatmaps below each pair display %CD69+ activation, R2 sequencing read counts, and enrichment scores for all tested variants.

**Figure S7.**
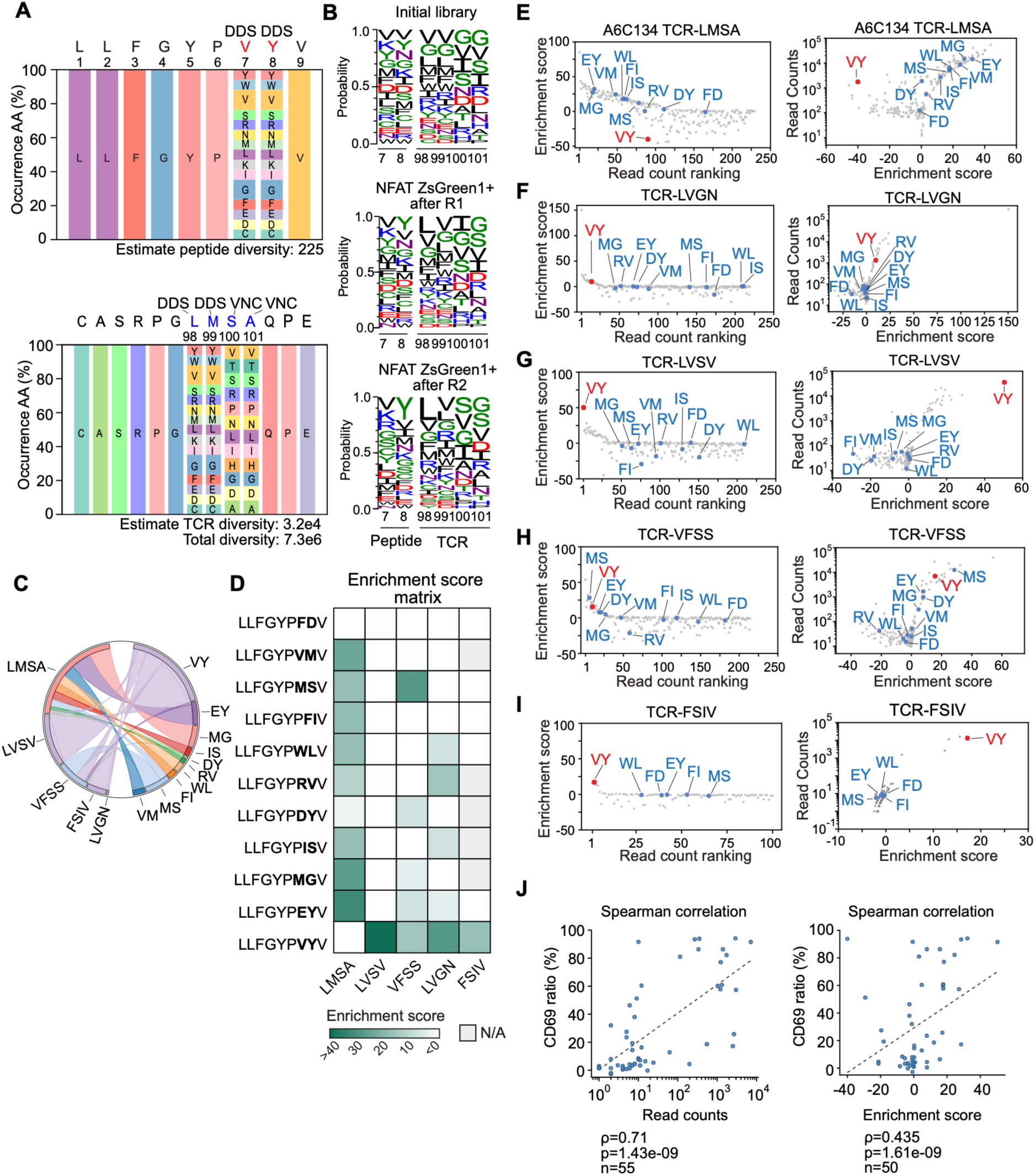
Library design and enrichment analysis for the A6C134–Tax library-on-library screen. Related to Figure 6. **(A)** Design for the A6C134–Tax co-evolutionary library. Position-frequency matrices showing amino acid diversity at each mutagenized position in the Tax peptide (left) and A6C134 CDR3β (right) libraries. Degenerate codons indicated above each position. Estimated library diversities are shown below. **(B)** Positional sequence logos for Tax peptide positions 7 and 8 and A6C134 CDR3β positions 98–101 in the initial library, after the first screening round, and after the second screening round, showing progressive convergence toward functional sequences. **(C)** Circos plot of experimentally validated TCR-peptides pairs. Ribbons connect each TCR clonotype (left) to its paired peptide partners (right), with ribbon width proportional to read count, illustrating the breadth and relative abundance of peptide partners recovered for each TCR clonotype. **(D)** Enrichment score heatmap of TCR-peptide variant matrix. Five TCR variants (LMSA, LVGN, LVSV, VFSS, FSIV) tested against 11 peptide variants. Bold residues indicate the variant positions on peptide sequence. Grey (N/A) indicates pairs with no detectable reads. **(E–I)** Scatter plots showing peptide variant partners for five TCR clonotypes, LMSA **(E)**, LVGN **(F)**, LVSV **(G)**, VFSS **(H)**, and FSIV **(I)**. Left plots show enrichment score (y-axis) plotted against read count ranking (x-axis); right plots show read count (y-axis) plotted against enrichment score (x-axis). The wild-type peptide VY is highlighted in red, other tested peptide variants are shown in blue. **(J)** Read counts and enrichment score correlate with T cell activation across validated TCR–pMHC pairs. Left plot shows read counts vs activation (ρ=0.71, p=1.43×10^-9^), Right plot shows correlation with enrichment score (ρ=0.435, p=1.61×10^-9^). Dashed lines indicate linear regression fits.

